# Retromer oligomerization drives SNX-BAR coat assembly and membrane constriction

**DOI:** 10.1101/2022.07.27.501715

**Authors:** Navin Gopaldass, Maria Giovanna De Leo, Thibault Courtellemont, Vincent Mercier, Christin Bissig, Aurélien Roux, Andreas Mayer

## Abstract

The retromer coat mediates protein exit from endosomes and impacts many signaling pathways, lysosomal biogenesis, and diseases such as Parkinson’s, Alzheimer’s and COVID-19. Retromer complexes (CSC in yeast) form coats by interconnecting sorting nexins (SNX). The dynamics of this process is poorly explored. Here, we analyze the oligomerization of CSC/SNX-BAR retromer coats on oriented synthetic lipid tubules. SNX-BARs and CSC assemble a static tubular coat that does not exchange subunits. Coat formation proceeds bidirectionally, adding new subunits at both ends of the coat. High concentrations of SNX-BARs alone suffice to constrict membrane tubes to an invariant radius of 19 nm. At lower concentrations, CSC-complexes must drive constriction, which requires their oligomerization. CSCs populate the SNX-BAR layer at densities that increase with the starting radius of the membrane tube. We hence propose that retromer-mediated crosslinking of SNX-BARs at variable densities tunes the coat according to the energy required to deform the membrane. This model is supported by the effects of mutations interfering with retromer oligomerization, which impair retromer function in yeast and human cells.

## Introduction

Endosomes and lysosomes form a complex network of interconnected organelles of different composition and function. They exchange proteins through fusion and fission with each other and, in a more selective fashion, through endosomal transport carriers (ETCs). ETCs are tubulo-vesicular structures bulging from the limiting membrane of these organelles. The formation of ETCs comprises several steps: Cargo selection, membrane deformation, and detachment from the donor organelle through membrane fission.

Alternatively, cargo can also pass between endo-lysosomal compartments through kiss-and-run, a transient fusion between two endo-lysosomal organelles followed by immediate re-fission (Solinger *et al*, 2020; Luzio *et al*, 2014). ETCs form through a variety protein coats and their interaction partners, such as the retromer, retriever and CCC complexes, ESCPE-1, sorting nexins and the WASH complex (Chen *et al*, 2019; Derivery *et al*, 2009; Gomez & Billadeau, 2009; Phillips-Krawczak *et al*, 2015; Rojas *et al*, 2007; Temkin *et al*, 2011; Lucas *et al*, 2016; Simonetti *et al*, 2022, 2019).

Retromer is a conserved tubular protein coat, which was originally defined in yeast as a stable complex that dissociates into two parts: The SNX complex, consisting of the SNX-BAR sorting nexins Vps5 and Vps17, and the peripheral CSC complex (Vps26, Vps29 and Vps35) (Seaman *et al*, 1998). In non-yeast systems, the Vps26/29/35 complex alone is referred to as retromer. It is recruited to membranes through various sorting nexins. Some of them carry BAR domains, such as the SNX-BARs Vps5 and Vps17, others do not, such as Snx3/Grd19 (Harrison *et al*, 2014; Deatherage *et al*, 2020; Strochlic *et al*, 2007; Harterink *et al*, 2011). The SNX complex (Vps5/Vps17) binds membranes via PX domains, which recognize phosphatidylinositol-3-phosphate (PI3P) (Burda *et al*, 2002), and through BAR domains, which bind highly curved membranes. The SNX complex recruits CSC, which by itself shows only weak affinity for the membrane, although it can interact with the bilayer when bound to other sorting nexins, such as Snx3 (Lucas *et al*, 2016; Purushothaman & Ungermann, 2018; Leneva *et al*, 2021; Strochlic *et al*, 2007; Deatherage *et al*, 2020). Retromer associates with numerous other factors, which are important for the formation of the transport carriers and/or their fission from the membrane. These include components of the Rab-GTPase system (Seaman *et al*, 2009; Rojas *et al*, 2008; Jia *et al*, 2016; Balderhaar *et al*, 2010; Liu *et al*, 2012; Purushothaman &

Ungermann, 2018), the actin-regulating WASH complex (Harbour *et al*, 2012; Chen *et al*, 2019; Derivery *et al*, 2009; Jia *et al*, 2012; Gomez & Billadeau, 2009; Phillips-Krawczak *et al*, 2015; Temkin *et al*, 2011; Lucas *et al*, 2016), or EHD1, an ATPase that has structural similarities to dynamins (Daumke *et al*, 2007; Gokool *et al*, 2007).

Structural studies of sorting nexins and retromer begin to elucidate how these coats wrap around membranes and how they recruit cargo (Leneva *et al*, 2021; Kovtun *et al*, 2018; Hierro *et al*, 2007; Lucas *et al*, 2016; Kendall *et al*, 2020; Collins *et al*, 2005, 2008; Purushothaman *et al*, 2017; Zhang *et al*, 2021). A further mechanistic analysis of the formation of ETCs and their fission from endo-lysosomal compartments will, however, require complementing dynamic data from in vitro systems that reproduce the formation and fission of ETCs in a well-defined, tunable and optically well-resolved setting. Attempts in this direction have already been undertaken. Retromer coat produces tubules on giant unilamellar vesicles (GUVs) (Purushothaman & Ungermann, 2018; Purushothaman *et al*, 2017). Those are hard to quantify because the tubules are numerous and difficult to resolve by light microscopy. Retromer oligomerization could also be followed on supported planar lipid bilayers (Deatherage *et al*, 2020), where protein interactions can be studied extremely well. But such a system appears less apt for observing tubulation by the coat and fission.

We engaged in an in vitro characterization of retromer because our studies on membrane fission on yeast vacuoles and mammalian endosomes revealed the PROPPINs Atg18 and WIPI1, respectively, as crucial factors (Zieger & Mayer, 2012; Peters *et al*, 2004; Gopaldass *et al*, 2017; DeLeo *et al*, 2021). Atg18 integrates with CSC to form the CROP complex (Courtellemont *et al*, 2022; Marquardt *et al*, 2022) displays much more potent membrane fission activity than the PROPPIN alone. To generate a system that may allow to analyse the mechanistic relationship of CROP to retromer, we used oriented lipid tubules on glass supports (Dar *et al*, 2015). The tubules allow to quantitatively follow the formation of the coat on them, a property which we exploited for an analysis of the properties and dynamics of retromer.

## Results

Supported membrane tubes (SMTs) (Dar *et al*, 2015) are individually observable tubular membranes that are immobilized and amenable to quantitative optical analysis. SMTs can be generated by liquid flow through a microfluidic chamber carrying lipid spots on its glass bottom. The flow produces arrays of parallel membrane tubes, which probably become stabilized in this orientation by occasional contacts with non-coated spots of the glass surface. Such SMTs allow to image the behaviour of proteins on the tubes over extended periods of time. The tube diameters can be quantified via a low percentage of incorporated fluorescent lipidic tracers because the fluorescence per unit tube length will depend on the number of fluorescent lipids in that unit, and hence upon the radius of the tube.

### CSC and SNX-BARs cooperate to constrict pre-formed membrane tubules to a uniform radius

We adapted this system to study the formation of retromer coats. To visualize SNX/CSC coat formation, an SMT array was formed on a coverslip that was covalently coated with polyethylene glycol and mounted in a flow chamber. The SMTs were labelled through 1 mol% of Texas Red DHPE and contained 5% PI3P, because this phosphoinositide is required to recruit SNX-BARs onto membranes (Yu & Lemmon, 2001; Cheever *et al*, 2001; Song *et al*, 2001). Upon addition of purified recombinant SNX complex (Vps5 and Vps17) and mClover-tagged CSC (Vps26, Vps35, Vps29^mClover^) (Suppl. Fig. 1a, b), uniform membrane binding could be observed by spinning disc fluorescence microscopy within seconds (Fig.1a, movie 1). PI3P and SNX were necessary to recruit CSC to the tubes (Suppl. Fig. 2a). About one minute after addition of SNX and CSC^mClover^, the mClover-signal strongly accumulated at multiple discrete sites on a tubule, suggesting that CSC was concentrating into separate protein domains (Fig. 1a, movie 1). The domains elongated over time, as visualized by kymograph analysis (Fig 1b). Co-labelling of CSC and SNX with mRuby and GFP, respectively, revealed that the zones where both CSC^mRuby^ and SNX^GFP^ were concentrated mirrored precisely zones of decreased lipid fluorescence, which were visualized through the red-fluorescing lipid Cy5.5-PE (Fig. 1c and d). The concentrations necessary for domain-formation by SNX complex and CSC varied from one preparation to another by a factor of two to three and were in general in the range of 10-25 nM. At such low concentrations, domains did not form in the absence of CSC (Fig. 1e).

**Figure 1:**
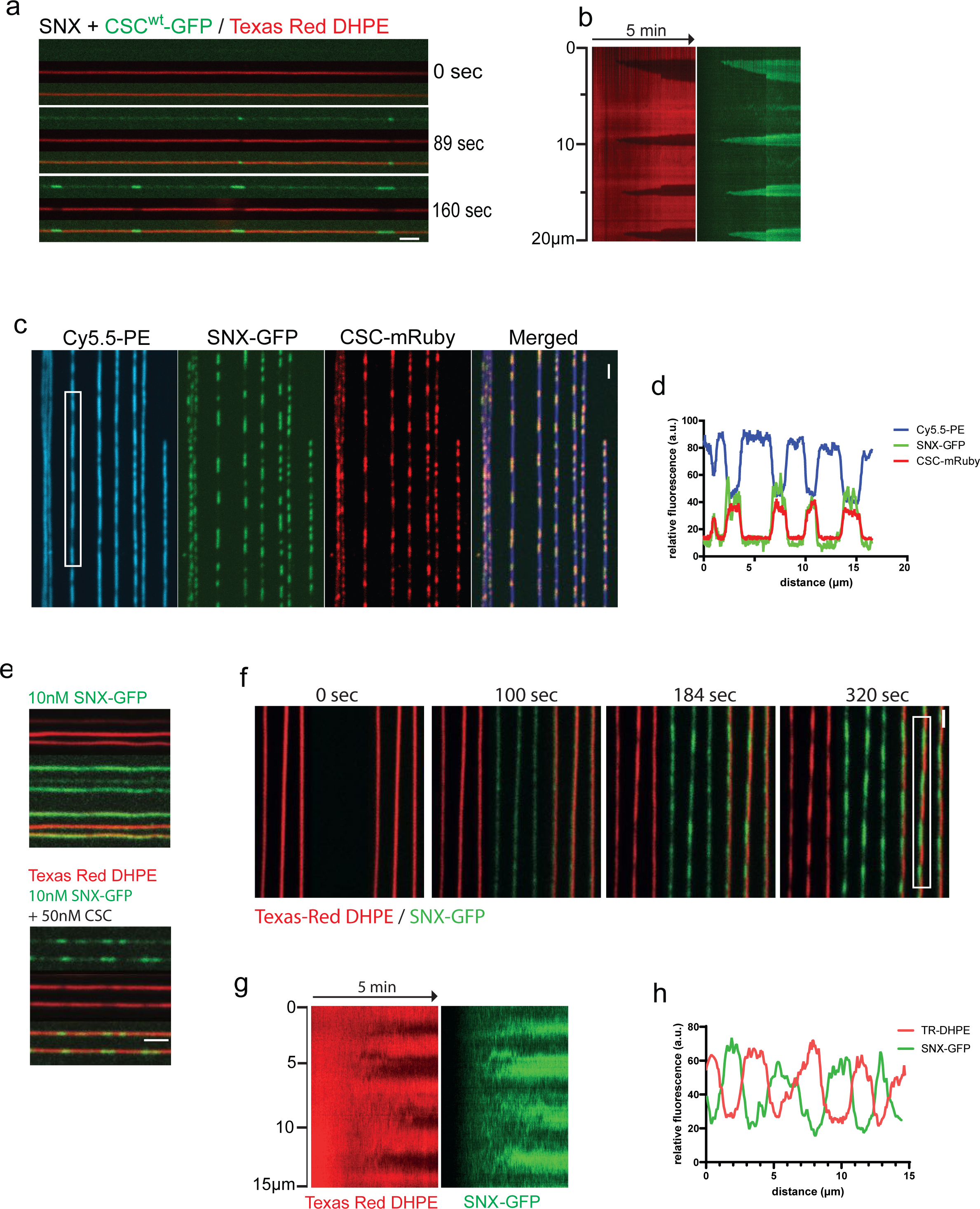
Assembly of retromer coats on supported membrane tubes. **a.** Dynamics of scaffold formation. 25 nM SNX and 25 nM CSC^mClover^ in PBS was added to SMTs and imaged by confocal microscopy at a frame rate of 1 Hz for 5 min. **b.** Kymograph of the tubule shown in a. See movie 1. **c.** SNX^GFP^ colocalizes with CSC^mRuby^ on SMTs. SMTs containing 1mol % of the fluorescent lipid Cy5.5-PE were incubated with 25 nM of SNX^GFP^ and CSC^mRuby^ for 2 min. Then, the tubes were imaged by confocal microscopy. **d.** Line scan analysis along the boxed tubule from c. **e.** Scaffold formation at low SNX concentration is facilitated by CSC. SMTs were incubated as in a, using 10 nM SNX^GFP^ in combination with either 50 nM CSC or only control buffer. **f.** Scaffold formation by elevated concentrations of SNX alone. 100 nM SNX-GFP was added to SMTs and imaged by confocal microscopy at a rate of 0.5 Hz for 5 min. **g.** Kymograph of the tubule highlighted in f. **h.** line scan analysis of the tubule highlighted in f. This experiment is also shown in movie 2.

Domains could also be formed by SNX alone (Fig. 1f, g and h, movie 2). However, for a given preparation this always required SNX concentrations 5-10 times above those that sufficed to generate domains in the presence of CSC.

The protein-enriched domains showed a strong reduction in lipid fluorescence, suggesting that the diameter of the underlying lipid bilayer was severely reduced (Fig. 1a-c, Suppl. Fig. 2b, c). The decrease in fluorescence was not due to a change in the direct environment of the fluorescent lipid upon protein binding (Jung *et al*, 2009; Hsieh *et al*, 2012), as we could observe the same effect using alternative lipid probes, which carry the fluorophore either on the membrane surface (Texas Red DHPE, Cy5.5-PE) or inside the hydrophobic part of the bilayer (NBD-PC) (Suppl. Fig. 2b; Fig. 1c, d). Therefore, we attribute the decrease in lipid fluorescence to a reduction in the amount of lipid underneath the protein domain, i.e., to a reduction in the radius of the membrane tube.

The radius of the membrane tubes in the constricted domains could be estimated using dynamin as a reference. Dynamin constricts membrane tubes to a defined radius of 11.2 nm (Roux *et al*, 2010). The lipid fluorescence in the SMTs can thus be calibrated and their radius can be deduced (Dar *et al*, 2015) (see Material and Methods and Supplementary Fig. 3 for details). This method revealed that the radius of membrane tubes underneath domains formed by CSC/SNX was 19.1 +/- 0.6 nm (Fig. 2 a-d), and that this radius was the same for membrane tubes constricted by high concentrations of SNX alone. Domains formed by SNX alone remained competent to bind CSC^mClover^ in a second incubation phase (Fig. 2e-f). However, the recruitment of CSC^mClover^ had no effect on lipid fluorescence under the pre-formed domains (Fig. 2g), suggesting that the tubes maintained their radius. This invariant radius was independent of the initial radius of the non-constricted tube, both for SNX/ CSC^mClover^ and for SNX-only domains (Fig. 2d). Thus, in agreement with structural studies (Kovtun *et al*, 2018; Leneva *et al*, 2021; Zhang *et al*, 2021; Kendall *et al*, 2020; Hierro *et al*, 2007; Lucas *et al*, 2016; Purushothaman *et al*, 2017), both CSC and SNX contribute to a constriction of the membrane tubes, probably by forming the retromer coat. SNX alone has membrane scaffolding activity, which defines the diameter of the coat independently of CSC, but CSC allows coat formation at lower SNX concentrations.

**Figure 2.**
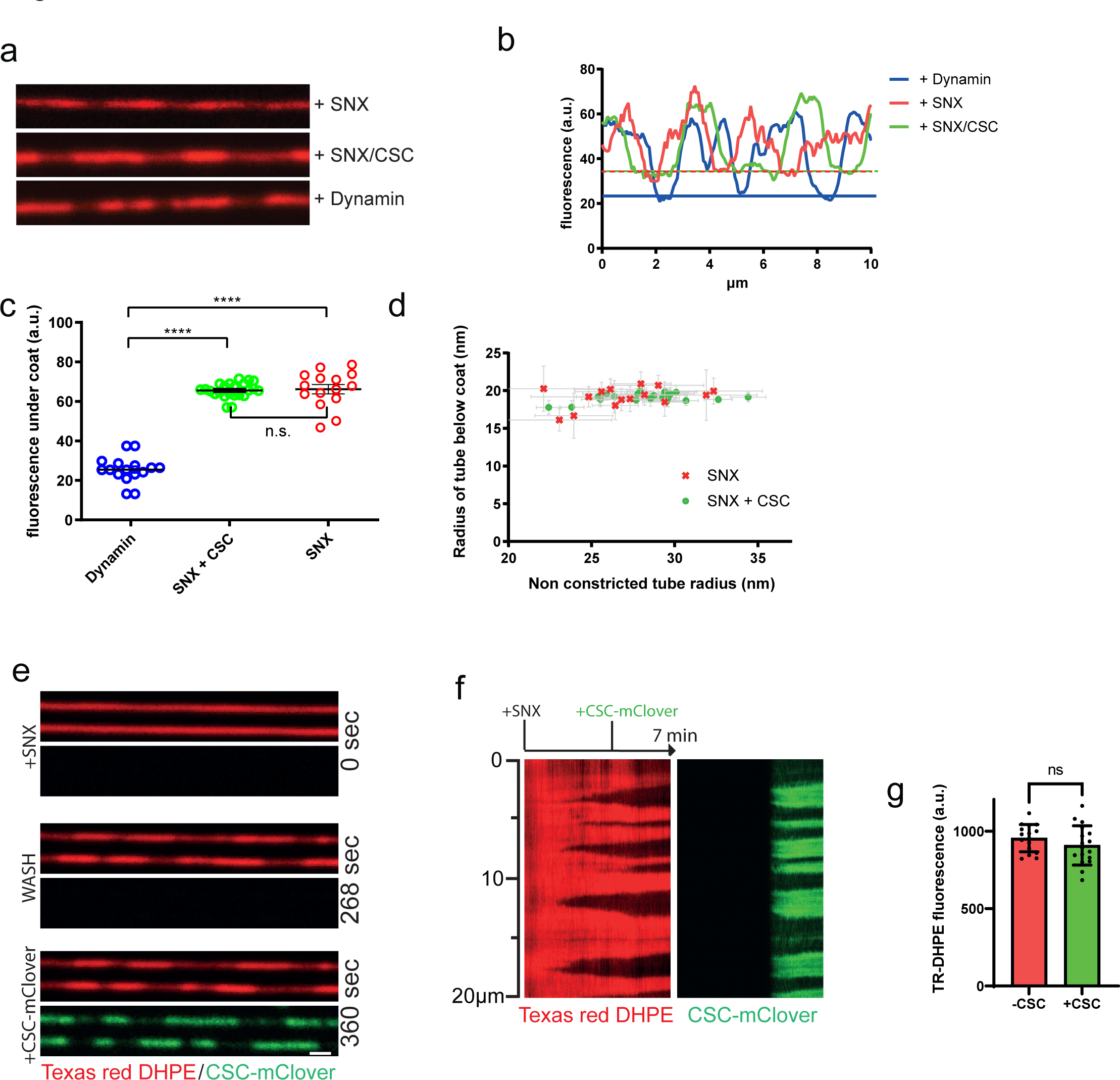
Constriction of membrane tubes by SNX and SNX/CSC. **a.** SMTs labelled with Texas-Red DHPE were incubated with non-tagged proteins at 25°C for 3 to 5 min and analyzed by fluorescence microscopy. Proteins were used at the following concentrations: 100 nM SNX; 25 nM SNX/25 nM CSC; 50 nM dynamin. **b.** Line scan analysis along the tubules from a. The lower boundaries of fluorescence are indicated by horizontal lines in the respective colors. **c.** Distribution of Texas-Red DHPE fluorescence in constricted domains for tubules coated by SNX (n = 16), SNX+CSC (n = 18), or dynamin (n = 15). ****: p < 0.0001 **d.** Constricted domain radius as a function of starting (non-constricted) tube radius. Radii of constricted and non-constricted regions of a variety of lipid tubes were determined using the known diameter of a dynamin-coated tube as a reference. Experiments were performed as in a, using 25 nM SNXs (n = 16) or 25 nM SNX plus 25 nM CSC (n = 18). **e.** Binding of CSC^mClover^ to constricted SNX domains. SMTs were first incubated with 100 nM SNX for 2 min at 25°C until constriction zones were visible through reduced lipid fluorescence. Then, 50 nM CSC^mClover^ was added under continuous acquisition at 0.5 Hz. **f.** Kymograph of a tubule from e. **g.** Quantification of Texas-Red-DHPE fluorescence under the constriction zone before and after CSC^mClover^ addition (n = 16). Error bars represent the standard deviation from the mean. n.s.: not significant (p=0.235).

### Retromer coats grow rapidly and bidirectionally

Cryo-EM analyses of retromer uncovered that the interface between the two Vps35 subunits of a retromer arch was asymmetric and that these subunits differed in overall structure (Leneva *et al*, 2021). It was proposed that this asymmetry might render coat assembly directional. Since the SMTs allow to observe coat growth in real time, we tested this hypothesis. To this end we first generated small red-fluorescing constricted coats using SNX and CSC^mRuby^ complexes. After a brief wash with protein-free buffer, SNX and CSC^mClover^ complexes were added (Fig. 3a to c). The green-fluorescing CSC^mClover^ extended the pre-existing red-fluorescing constricted zones that had been formed by CSC^mRuby^. The extension speed of the coat was substantial, ranging from 1 µm/min to 1.5 µm/min. This is in a similar range as the speed of dynamin polymerization (Roux *et al*, 2010). The elongation speed of this coat can also be put into perspective by comparison to the speeds of polymerization of actin tails (up to 3.6 µm/min; (Cameron *et al*, 1999)), or microtubules (10 µm/sec; (Gierke *et al*, 2010)). Based on the proposed structure of the retromer coat (Leneva *et al*, 2021; Kovtun *et al*, 2018) we estimate the observed extension speed to require the addition of approximately 10 to 15 SNX dimers per second at each end. CSC^mClover^/SNX elongated the pre-existing red-fluorescing coats at both ends with similar rates (Fig. 3b and c). Thus, despite the asymmetry in the arches (Leneva *et al*, 2021), the retromer coat displayed no inherent directionality of growth.

**Fig. 3:**
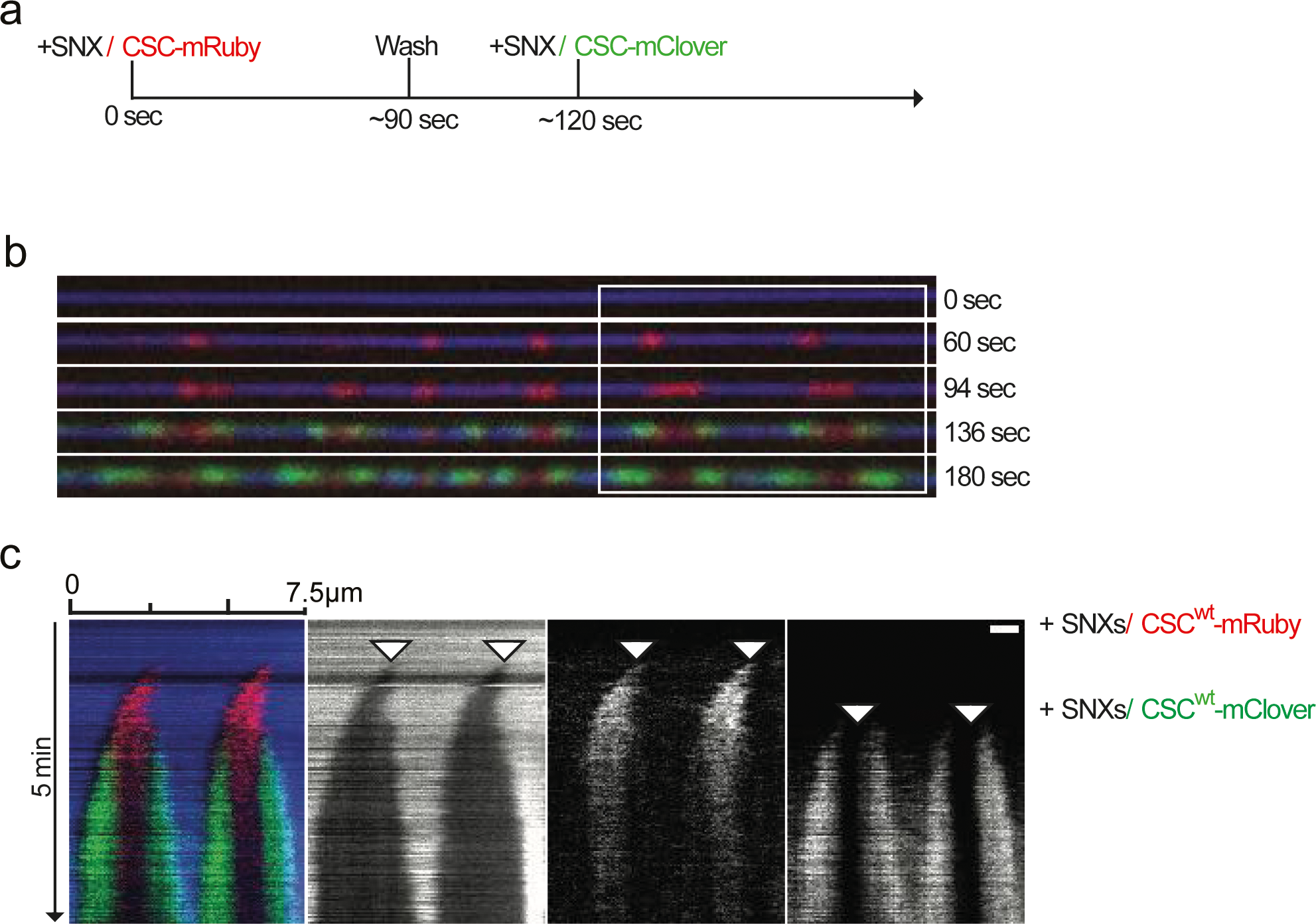
Bidirectional elongation of the coat. **a.** Scheme of the experiment **b.** SMTs labelled with Cy5.5-PE were first incubated with 50 nM SNX and 50 nM CSC^mRuby^ at 25°C until coat formation was initiated (∼90 sec). Then, non-bound SNX and CSC^mRuby^ were washed out and the tubes were subjected to a second incubation with 50 nM SNX and 50 nM CSC^mClover^ (3 min). Tubes were imaged by confocal microscopy at a framerate of 0.5 Hz. **c**. Kymograph of the tubule shown in b.

### The retromer coat is a static structure that does not exchange subunits

Structural studies of retromer-coated tubules formed with Vps5 revealed that these retromer coats are irregular in terms of their coverage with protein and that CSC oligomerizes into arch-like structures on the sorting nexins (Leneva *et al*, 2021; Kovtun *et al*, 2018). The irregularity raises several questions: Does the tubular coat represent a static structure, or is it rather dynamic, with subunits readily moving in and out? Does CSC facilitate SNX coat formation by dimerization and does the apparent variability in the occupancy of the SNX layer by CSC have functional implications? The formation of a static structure by SNX/CSC implies that the proteins form a rigid coat that might stabilize the underlying membrane tubule. To test this, we used an assay where a membrane tubule is pulled out of a giant unilamellar vesicle (GUV) by means of an optical trap. Through analysing the displacement of the bead in the trap, such a setup allows direct measurement of the force required to generate and maintain the tubule (Roux *et al*, 2010). Shortly after SNX/CSC^mClover^ addition, protein bound the pulled tubule (Fig. 4a). As a consequence, the pulling force exerted on the bead decreased sharply (Fig 4b). Elongating this tubule by displacing the GUV transiently increased in the force again (Fig. 4c and d), but as the SNX/CSC^mClover^ coat grew along the newly extracted portion of the tubule (Fig. 4e), the force decreased again. This could be repeated several times (Fig. 4d). Together, these data suggest that the SNX/CSC coat forms a rigid scaffold that suffices to stabilizes a membrane tubule.

**Fig. 4:**
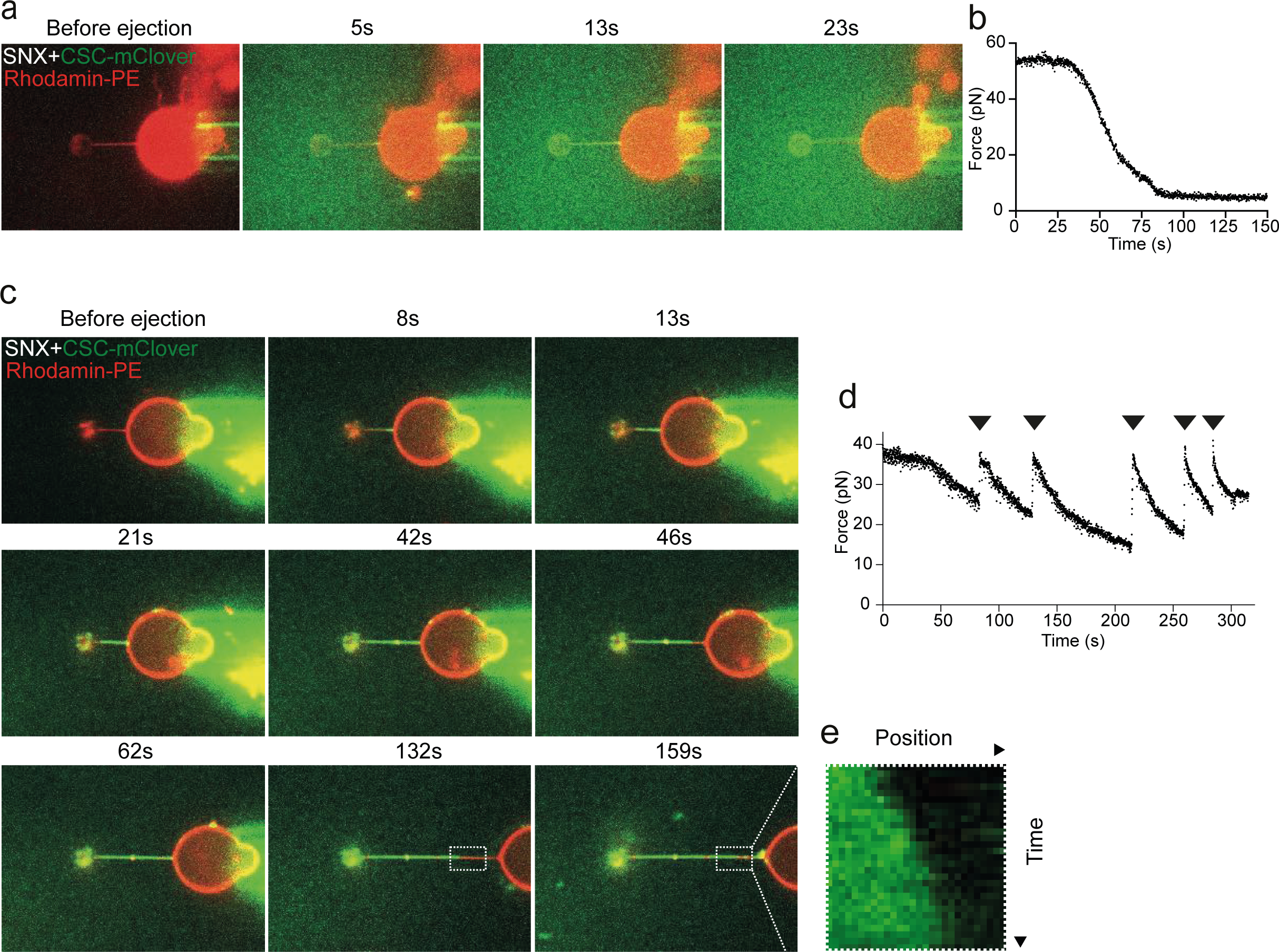
Stabilization of pulled membrane tubes by CSC/SNX. **a.** Coat formation on a pulled membrane tubule. Confocal pictures of a GUV labelled with Rhodamine-DHPE (red). A membrane tubule has been pulled from the GUV through a small bead and optical tweezers. The GUV is shown before and at several time points after ejection of SNX/CSC^mClover^ (green) from a pipette in the vicinity of the GUV. **b.** Measurement of the force exerted on the bead as a function of time after protein ejection, taken from the experiment in a. **c.** Repetitive pulling and stabilization. Confocal pictures of a GUV labelled with Rhodamine-DHPE (red). A tubule has been pulled as in a and SNX/CSC^mClover^ (green) was added. The GUV is shown before and after protein ejection, and at several stages of subsequent re-pulling and stabilization through additional coat recruitment. Protein quickly populates new tube regions generated by pulling back the GUV. **d.** Measurement of the force exerted on the bead as a function of time for the experiment shown in c. Arrowheads mark the timepoints when the GUV has been pulled back. **e.** Kymograph of the portion of the tubule boxed in d, showing growth of retromer coat into a newly pulled portion of the tubule.

### CSC density may define the work that the coat performs in deforming membranes

Binding of CSC adds further interactions to the SNX layer (Kovtun *et al*, 2018; Leneva *et al*, 2021; Lucas *et al*, 2016), and could hence provide additional energy for membrane scaffolding. This led us to test whether coat stoichiometry might vary as a function of the radius of the starting tubule, because constricting a less curved membrane requires more work (Roux, 2013). We first assayed whether the coats are saturated, using SNX and CSC separately in a two-stage experiment. Constricted coats on SMTs were pre-formed from red-fluorescing SNX/CSC^mRuby^ complexes. Non-bound proteins were washed away and, in a second incubation, we added either green-fluorescing SNX^GFP^ or CSC^mClover^ (Fig. 5 a, b). SNX^GFP^ was recruited to the non-constricted areas of the tubes but could not integrate into the constricted domains, suggesting that the membrane in these domains was fully covered. By contrast, CSC^mClover^ bound mainly to the constricted domains. CSC^mClover^ was not recruited in exchange for pre-bound CSC^mRuby^, which might have dissociated from the constriction, because the CSC^mRuby^ signal in the constricted domains remained constant after the addition of CSC^mClover^ (Fig. 5c). This suggests that the constricted domains are saturated for SNX complexes but retain free binding sites for additional CSC.

**Figure 5:**
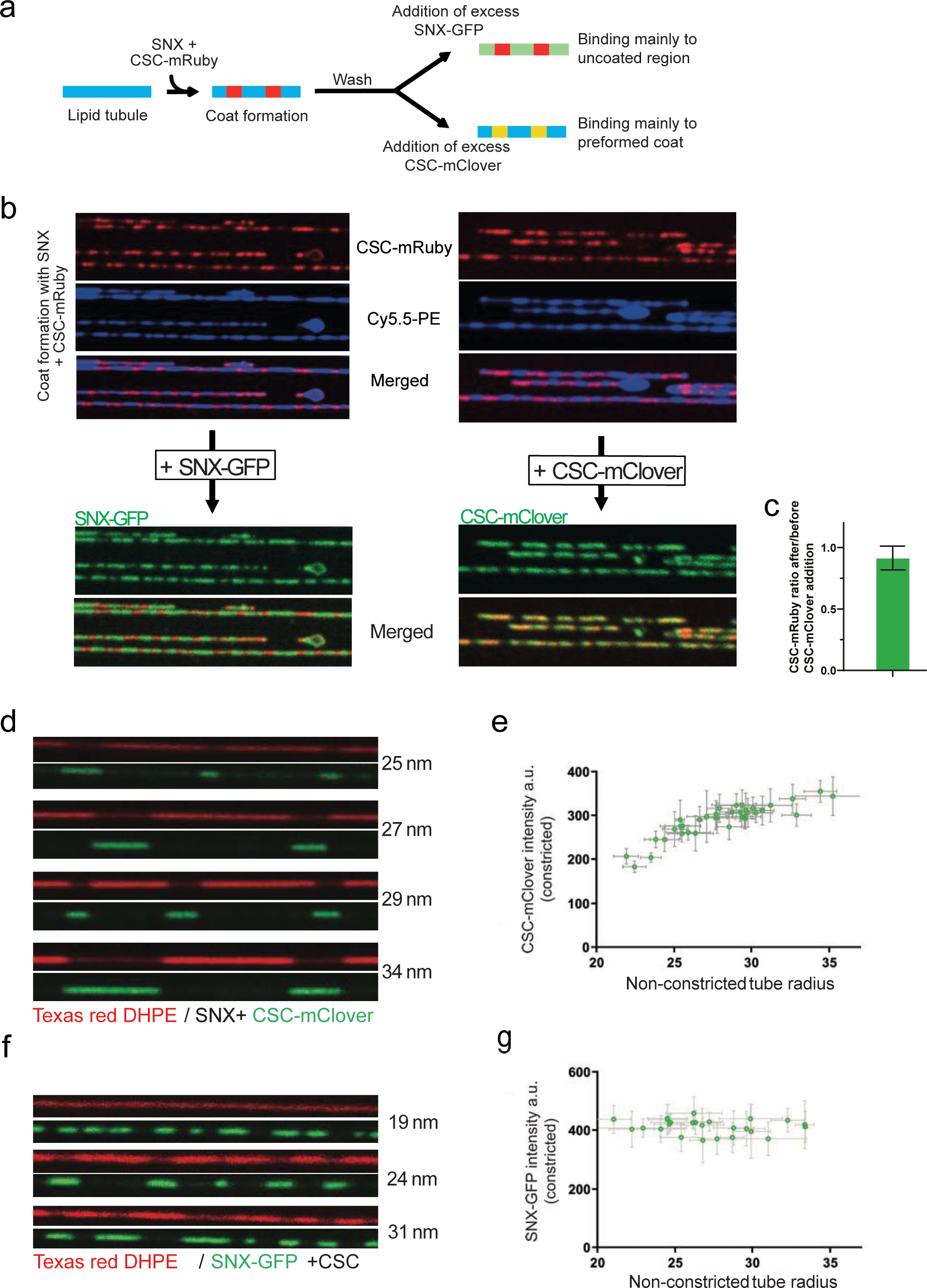
Variable saturation of the SNX layer with CSC. **a.** Recruitment of additional subunits to pre-formed coats using differentially labelled CSC and SNX. Scheme of the experiment shown in b and c. **b.** Coats were formed on SMTs using 25 nM SNX and 25 nm CSC^mRuby^. Excess protein was washed out with buffer and 50 nM SNX^GFP^ or 50 nM CSC^mClover^ were added. SMTs were imaged after SNX/CSC^mRuby^ coat formation and 2 min after addition of CSC^mClover^ or SNX^GFP^. **c.** Ratio of CSC^mRuby^ signals in the constricted areas before and after addition of CSC^mClo-ver.^ **d.** Occupation of SNX domains with CSC as a function of the starting radius of the tube (naked tube radius). Arrays of SMTs were incubated with 25 nM SNX and 25 nM CSC^mClo-ver^. The density of CSC^mClover^ in constricted domains was traced through its fluorescence signal. The starting radius of the tube was estimated through Texas Red-DHPE fluorescence in non-constricted regions and calibration with dynamin. This radius is indicated for each tube. **e.** The density of CSC^mClover^ in SNX/CSC^mClover^ coats from d was plotted as a function of the radius of the non-constricted tube. **f.** Experiment as in d using 25 nM SNX^GFP^ and 25 nM CSC. **g.** The fluorescence signals of SNX^GFP^ in SNX/CSC coats from f were quantified plotted as a function of the radius of the non-constricted tube as in e.

To test whether the density of CSC on retromer coats has functional significance, we compared the density of CSC in constricted coats that had had been formed on membrane tubes of different starting radius. The SMT system is very apt for this analysis because it simultaneously generates many tubes of variable radii on the same slide. Calibration via the integrated fluorescent lipids showed that these “naked tube” radii varied mostly from 20-40 nm under the conditions we employed. We measured the radius of non-constricted regions to approximate the starting radius of the tube and then measured the signals of SNX^GFP^ and CSC^mClover^ in the constricted domains of that tube (Suppl. Fig. 3). Whereas CSC^mClover^ fluorescence per unit length of constricted tube increased as a function of starting tube radius (Fig. 5d, e), the density of SNX^GFP^ in the constricted domains was independent of the starting tube radius (Fig. 5f, g). Thus, the larger the starting tube, the more CSC is incorporated into the constricted coat. Since the energy required to constrict a membrane tube to a defined diameter increases with its initial radius, this suggests that the work that the SNX coat performs to constrict the membrane may be tuned through the density of CSC complexes that are incorporated to connect them.

### Retromer oligomerization supports coat constriction

CSC arches can connect multiple SNX subunits and can form oligomers (Deatherage *et al*, 2020; Kendall *et al*, 2020; Kovtun *et al*, 2018; Lucas *et al*, 2016). Oligomerization might provide additional bonds for membrane deformation by the coat and/or facilitate coat assembly by constraining the subunits in an orientation relative to each other that is best compatible with a constricted lipid tube. In both cases, the capacity of CSC to oligomerize should play an important role for driving the formation of constricted domains. Structural studies showed that CSC forms dimers through a conserved interface on Vps35 (Kendall *et al*, 2020; Leneva *et al*, 2021). To assess the contribution of retromer dimerization on coat formation we used the PDB-PISA software (https://www.ebi.ac.uk/pdbe/pisa/) to model the Vps35-Vps35 dimerization interface, using a retromer CryoEM structure (Kovtun *et al*, 2018; Leneva *et al*, 2021) as an input. PDB-PISA calculates the energy contribution of each residue to a protein-protein interaction surface. This approach predicts residues to form hydrogen bonds or salt bridges between the two Vps35 subunits. We selected the conserved Vps35 residues D671, L722 and R775 for substitution by alanine, yielding the *vps35^PISA^* allele (Fig. 6a, b). We also generated *vps35^AAA3KE^*, in which another set of conserved residues in the interaction region are substituted. The AAA3KE substitutions abolish the capacity of mammalian VPS35 to self-associate and lead to partial secretion of the vacuolar protease CPY in yeast (Kendall *et al*, 2020). A recent structural study showed that all these substituted residues contribute to an asymmetric Vps35-Vps35 interface (Leneva *et al*, 2021). We extracted CSC complexes containing both Vps35 variants from yeast (Fig. 6c) and tested their capacity to form higher order oligomers by blue native gel electrophoresis (Fig. 6d). CSC^wt^ migrated in three main bands at apparent molecular masses compatible with a Vps29^mClover^-containing monomer of CSC (207 kDa), a dimer (414 kDa), and a tetramer (828 kDa). The most slowly migrating species was abolished in CSC from *vps35^AAA3KE^* cells and weaker in CSC from *vps35^PISA^*, while the intermediate-sized forms persisted. This suggests that the slowest form represents a CSC tetramer held together by Vps35 dimerization, whereas the dimer may persist through a Vps26-Vps26 interface (Kovtun *et al*, 2018).

**Figure 6.**
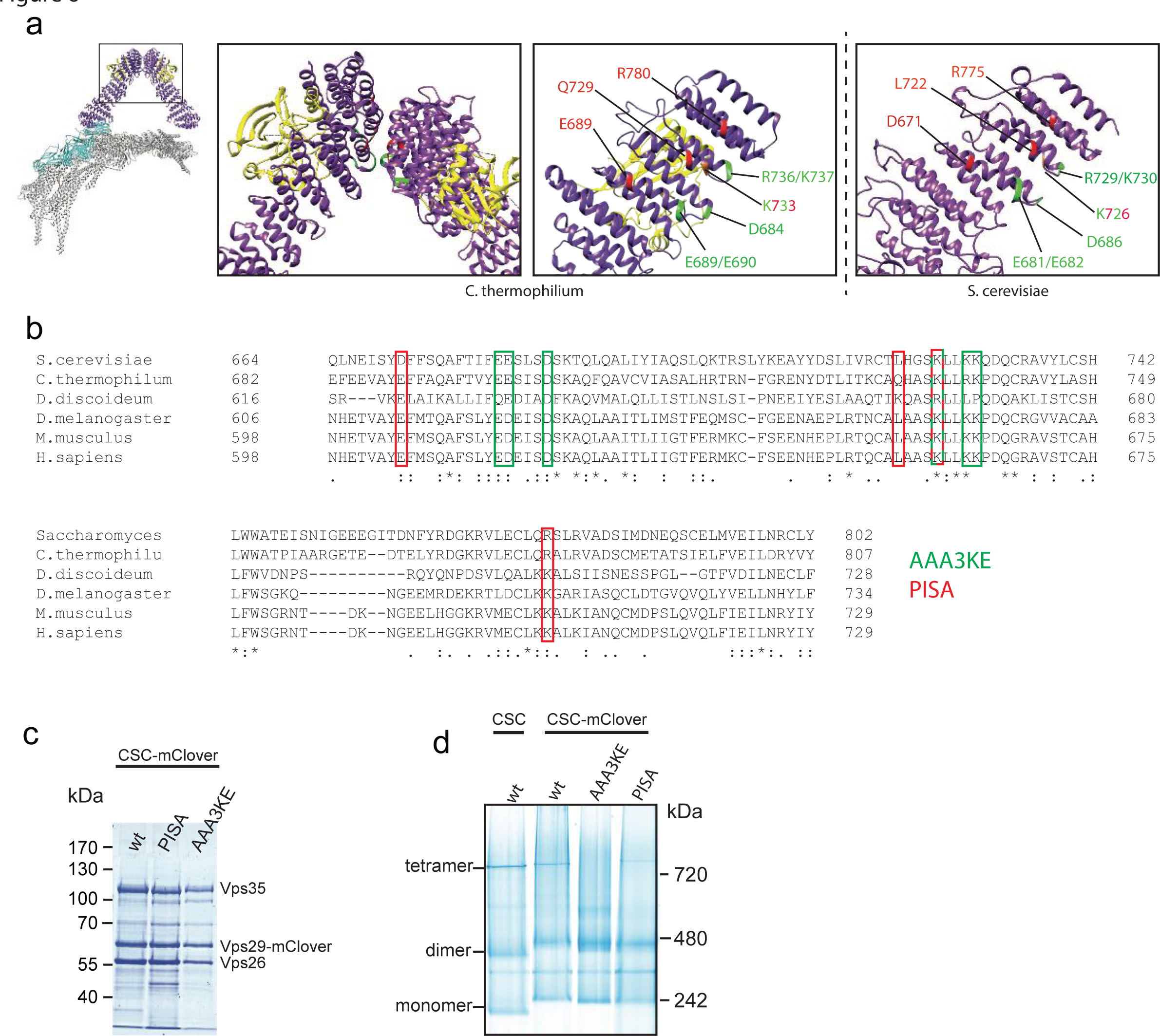
Substitutions destabilizing the Vps35-Vps35 interface. **a.** Structure of the pentameric retromer complex (Leneva *et al*, 2021). The boxes highlight the Vps35 dimerization interface. Residues substituted in *vps35^PISA^* and *vps35^AAA3KE^*are shown red and green, respectively, in the structure from C. thermophilum and in a model of the S. cerevisiae complex derived from the Chaetomium thermophilum structure (PDB 7BLR using the online modelling tool Swiss-model (https://swissmodel.expasy.org). **b.** Sequence alignment of the Vps35 dimerization domains from different species. Amino acids substituted in *vps35^PISA^*and *vps35^AAA3KE^* are shown in red and green, respectively. One residue (in red-green) is shared between the two. **c.** Coomassie-stained SDS-PAGE gel of purified CSC^mClover^ complexes containing the indicated Vps35 variants. **d.** Blue native PAGE gel showing the formation of higher order assemblies for CSC^mClover^ complexes containing Vps35 variants and their tentative assignment as monomers, dimers and tetramers.

We performed SMT assays to assess the capacity of both CSC variants to form constricted domains. To avoid potential influences of the CSC variants on the speed or extent of SNX recruitment to the tubes, the experiments were performed in two phases. A first incubation at low SNX concentration (25 nM) allowed this complex to bind the tubes without forming constricted domains (Fig. 7a, b). Unbound SNX was washed away before CSC^mClover^ variants were added for the second incubation phase. CSC^mClover^ with *vps35^AAA3KE^* and *vps35^PISA^* was recruited to the prebound SNX with similar kinetics and to similar extent as the wildtype complex (Fig. 7b-d). However, only wildtype CSC drove the formation of constricted domains (Fig. 7 b, c). The CSC variants also failed to drive constriction when they were co-incubated with SNX right from the beginning in a one-phase experiment (Suppl. Fig. 4a-d). That the CSC variants were in principle able to bind a constricted SNX layer was shown by a further experiment, in which constricted SNX-only coats were pre-formed in a first incubation phase at high SNX concentration. CSC*^AAA3KE-mClover^* and CSC^PISA-mClover^ bound to those preformed constrictions similarly as the wildtype complex (Suppl. Fig. 4 e, f). Together, these results suggest that higher order self-assembly of CSC via the conserved Vps35 interface is necessary to drive membrane constriction by the SNX/CSC coat.

**Figure 7:**
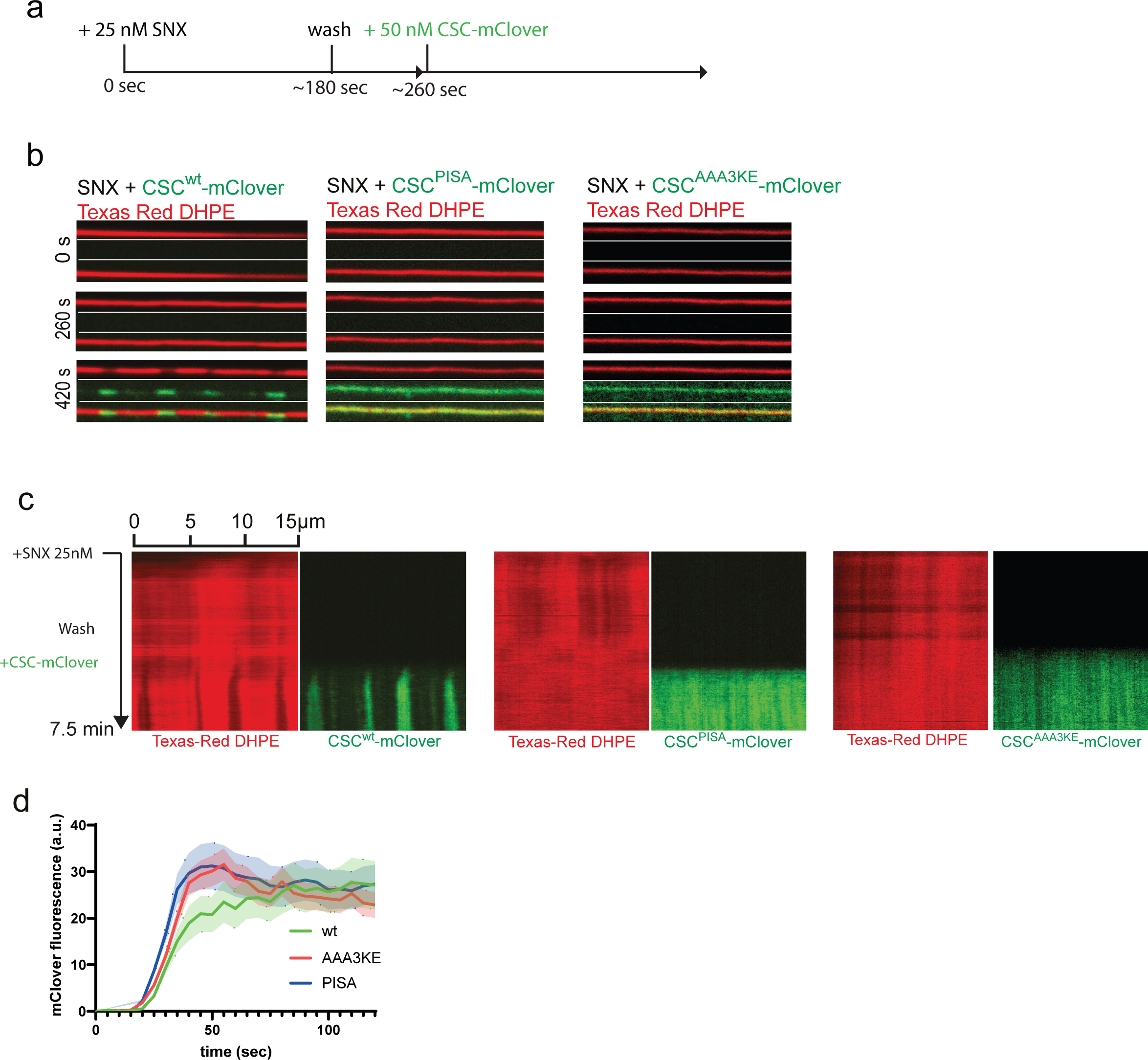
Effect of Vps35 dimerization on coat constriction. **a.** Experimental setup: SMTs labelled with Texas Red DHPE were preincubated with 25 nM SNX for 3 min to load them with SNX but not allow formation of constrictions. After a wash with protein-free buffer, 50 nM of CSC^mClover^ carrying the indicated Vps35 variants was added. The tubes were imaged by confocal microscopy at a framerate of 0.5 Hz. **b** Images of three timepoints after addition of CSC^mClover^ variants. **c** Kymographs of the entire reactions. Experiments are shown in movies 3-5. **d**. Recruitment kinetics of CSC^mClover^ variants to SNX-decorated SMTs during the second incubation phase. mClover fluorescence appearing along the entire length of the tubes was quantified over time. n=10 tubes per variant. Curves represent the mean and shaded areas around the curves represent the SEM.

### Mutations affecting retromer oligomerization impair cargo sorting in vivo

We used *vps35^AAA3KE^* and *vps35^PISA^*to test the relevance of CSC oligomerization in vivo. To this end, VPS35 was TAP-tagged and corresponding nucleotide exchanges were made at the VPS35 genomic locus, making the mutated alleles the sole source of Vps35 protein. Both *vps35^AAA3KE^* and *vps35^PISA^* were expressed at similar levels as a *VPS35^WT^*allele. (Suppl. Fig. 5a). They supported normal localization and abundance of yomCherry fusions of the CSC subunit Vps29 and the SNX subunit Vps17, suggesting that they are folded (Suppl. Fig. 5b and c). In contrast to an earlier study, which used secretion of the vacuolar protease CPY as an indirect assay and found only a very mild impact of *vps35^AAA3KE^* (Kendall *et al*, 2020), we assessed retromer function through microscopic localization of a yEGFP fusion of Vps10. Vps10 is a cargo receptor that uses retromer for returning from the pre-vacuolar compartment (the equivalent of a late endosome) to the trans-Golgi network (TGN) (Marcusson *et al*, 1994). Cells expressing wildtype VPS35 showed Vps10^yEGFP^ mostly in small dots scattered in the cytosol or adjacent to the vacuole (Fig. 8a), consistent with its expected location in the TGN and pre-vacuolar compartment (Chi *et al*, 2014). By contrast, cells lacking VPS35 (*vps35Δ*) accumulated significant amounts of Vps10^yEGFP^ on the vacuolar membrane, where they co-localized with the lipidic vacuole stain FM4-64 (Fig. 8a, b). This vacuolar localization is a hallmark of defective retromer function in yeast. It results from the failure to recycle Vps10 from the pre-vacuolar compartment before it finally fuses with the vacuole. The *vps35^AAA3KE^* and *vps35^PISA^* alleles produced an intermediate phenotype, where vacuoles were significantly more labelled by Vps10^yEGFP^ than in wildtype, but less than in *vps35Δ* (Fig. 8 a and b). The pre-vacuolar compartment recruits SNX and CSC (Burda *et al*, 2002; Liu *et al*, 2012). In line with this, Vps17^yomCherry^ or Vps29^yomCherry^ appeared as scattered dots when visualized by fluorescence microscopy (Suppl. Fig. 5b, c). We quantified the number of Vps10^yEGFP^ dots that were also Vps17^yomCherry^ positive. While 50% of Vps10^yEGFP^ positive dots in wildtype cells co-localized with Vps17^yomCherry^, 80-90% of colocalization was observed in cells expressing *vps35^AAA3KE^*or *vps35^PISA^* (Suppl. Fig. 5d). This is consistent with Vps10 being collected into SNX-containing structures but unable to recycle back to the Golgi.

**Figure 8:**
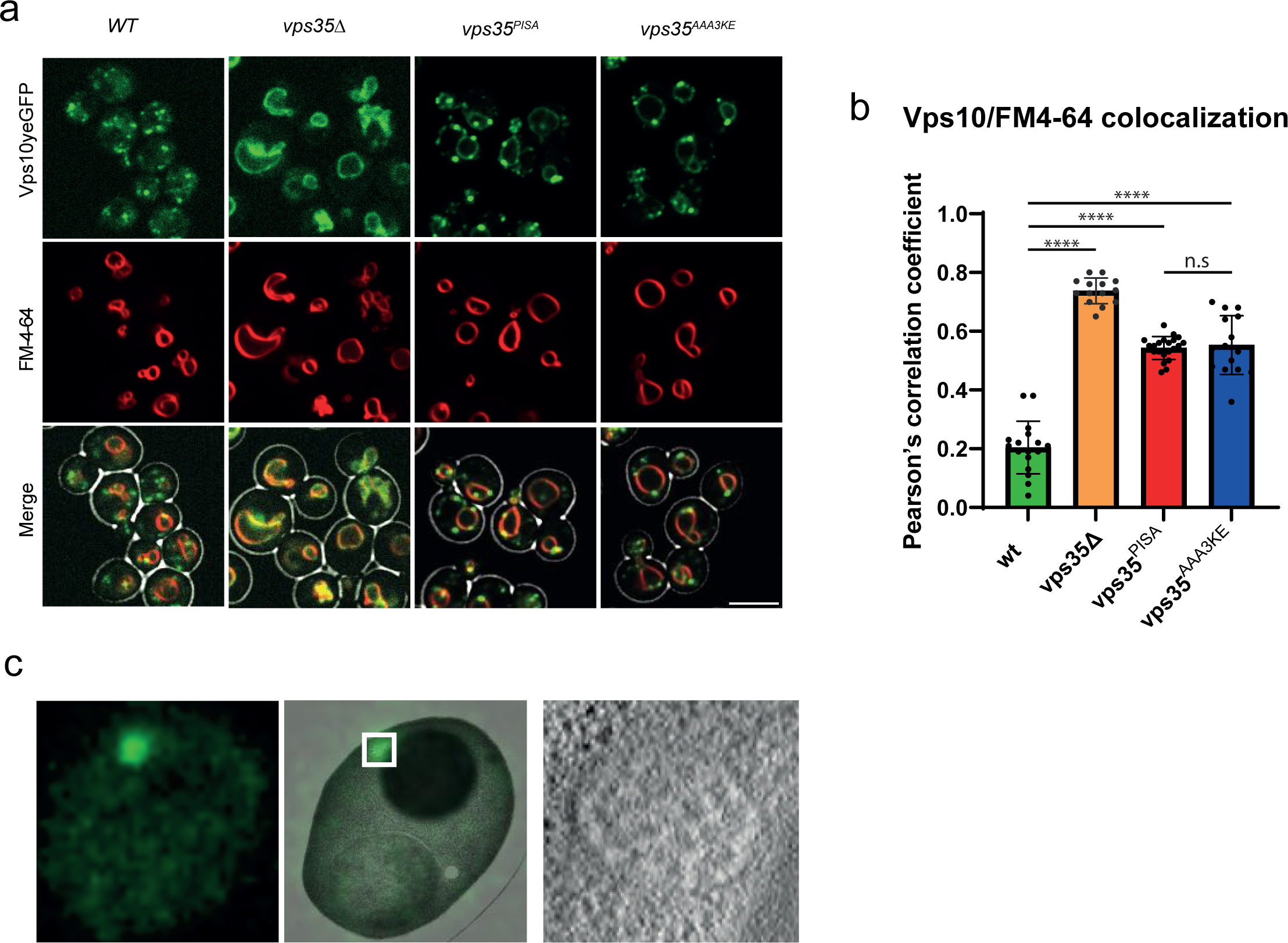
In vivo effect of Vps35 dimerization mutants on Vps10 in yeast. **a.** Vps10^yEGFP^ localization. Yeast cells carrying Vps10^yEGFP^ and expressing the indicated vps35 alleles as the sole source of Vps35 were logarithmically grown in SC medium. Their vacuoles were labelled with FM4-64. Cells were harvested by brief centrifugation and immediately imaged by confocal microscopy. Single confocal planes are shown. A brightfield image was used to outline the cell boundaries (shown in the merged images). Scale bar: 5 µM. **b.** Co-localization of Vps10^yEGFP^ and FM4-64 in cells from a was measured using Pearson’s coefficient. **c.** CLEM analysis of Vps10^yEGFP^ localization in vps35^PISA^ mutant cells. Logarithmically growing cells were high-pressure frozen and processed by freeze substitution and embedding.

Correlative light and electron microscopy of the Vps10^yEGFP^ dots in *vps35^PISA^* cells showed that the structures accumulating Vps17 were indeed PVCs, because they carried multiple lumenal vesicles, which is characteristic for these compartments (Fig. 8c).

As a readout for retromer function in human cells, we used the glucose transporter GLUT1, a well-characterized retromer cargo (Liu *et al*, 2012; Steinberg *et al*, 2013; Hesketh *et al*, 2014; Kvainickas *et al*, 2017; Evans *et al*, 2020). GLUT1 is normally localized at the plasma membrane and this localization requires its retromer-dependent export from endosomes. Knock-down of hVPS35 in HK2 cells resulted in a strong reduction of GLUT1 on the plasma membrane, consistent with a lack of its retromerdependent recycling (Fig. 9a). Expression of an si-RNA resistant form of hVPS35 rescued this phenotype, while expression of the corresponding mammalian mutant alleles hVPS35^AAA3KE^ and hVPS35^PISA^ did not (Fig. 9b, c). Expressing the AAAKE and PISA variants in wildtype cells led to a similar GLUT1 recycling phenotype as in the knock-down cells, while expression of the wildtype protein did not (Suppl. Fig. 6). This dominant negative effect suggests that the mutant proteins are correctly folded such that they can compete with the endogenous wt version for retromer complex formation. Another striking phenotype of cells lacking hVPS35 is an increase in the size of lysosomal compartments, probably due to a lack of membrane recycling and/or accumulation of undigested material resulting from insufficient delivery of lysosomal enzymes (Cui *et al*, 2018). hVps35 knockdown cells showed bigger LAMP1-positive lysosomal compartments (Suppl. Fig. 7a).

**Fig. 9:**
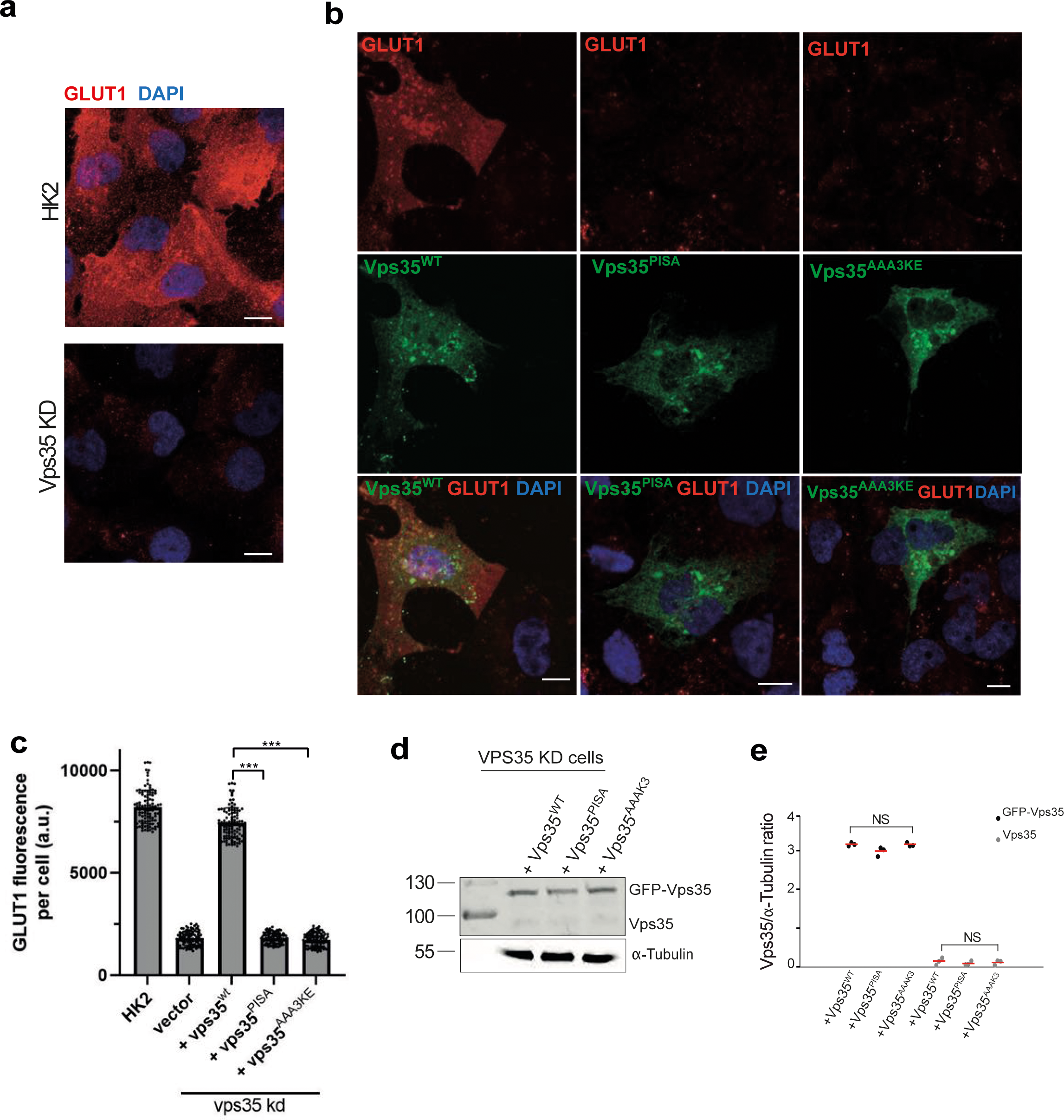
Effects of Vps35 dimerization mutants in human kidney (HK2) cells. **a.** GLUT1 at the plasma membrane. HK2 cells were treated with siRNA targeting VPS35 or with mock siRNA. Cells were fixed and stained with antibody to GLUT1 (red) and with DAPI (blue). Cells were not detergent permeabilized in order to preferentially show GLUT1 at the cell surface. Maximum projections of image stacks (step size in z of 300 nm) are shown. Scale bars: 10 μm. **b.** Influence of VPS35 variants on GLUT1. HK2 cells silenced for VPS35 were transfected with a plasmid carrying siRNA resistant wildtype or mutant forms of GFP-VPS35. GLUT1 was detected by fixation and immunofluorescence staining as in a. Maximum projections of image stacks (step size in z of 300 nm) are shown. Scale bars: 10 µm. **c.** Quantification of GLUT1 immunofluorescence in cells from (b). Regions of interest (ROIs) corresponding to cells expressing the indicated VPS35 variants, and some regions outside the cells (background), were manually defined using ImageJ software. Total cell fluorescence was integrated and corrected for background fluorescence. 105 cells per condition stemming from 3 independent experiments were analyzed. P values were calculated by Welch’s t-test. The analysis was performed with 99% confidence: *** p < 0.001. **d.** Expression of Vps35 variants in cells from b was analyzed by SDS-PAGE and Western blot against Vps35. Tubulin served as loading control. **e.** Quantification of the Vps35/tubulin ratio in cells from b. Data stems from three independent experiments and show the mean. NS: not significant (p>0.01).

These enlarged compartments could be brought back to normal size by expressing an siRNA resistant form of hVPS35, while expression of hVPS35^PISA^ and hVPS35^AAA3KE^ failed to rescue this phenotype (Suppl. Fig. 7b). Altogether, these data suggest a conserved role for Vps35 oligomerization in both yeast and human cells.

## Discussion

Structural analyses have revealed many important features of retromer coats (Hierro *et al*, 2007; Lucas *et al*, 2016; Kendall *et al*, 2020; Collins *et al*, 2005, 2008; Zhang *et al*, 2021; Purushothaman *et al*, 2017). Both Snx3- and Vps5-based coats show CSC forming arch-like structures that interconnect the sorting nexins over more than 20 nm and angular sections of around 60°C (Leneva *et al*, 2021; Kovtun *et al*, 2018). These models show limited regularity of the coat, both with respect to the placement of sorting nexins and their coverage with CSC (Leneva *et al*, 2021; Kovtun *et al*, 2018). This irregularity was suggested to represent potential plasticity that may, for example, allow the coat to adjust to different membrane curvatures or to integrate other proteins. Dynamic aspects of coat assembly have, however, not yet been experimentally tested. Our analyses of retromer coat formation in real time provide complementing functional information that relates to several features of the structural models. In our experiments with supported membrane tubes, SNX/CSC assembled into a coat that constricted membranes of variable starting curvature to an invariant radius of 19 nm. This number, obtained with a coat including Vps5 and Vps17, is in the range of the radius of 15 nm that was obtained in a structural study of a CSC coat formed with Vps5 alone (Kovtun *et al*, 2018) and similar to the radius of tubules formed by mammalian Vps5 homologs (Weering *et al*, 2012). Thus, even though both yeast SNX-BAR proteins, Vps5 and Vps17, are required for retromer function (Horazdovsky *et al*, 1997; Seaman & Williams, 2002), absence of one of them does not have a major impact on the dimensions of the membrane tubules shaped by retromer.

The coat formed by SNX/CSC appears as a stable and static scaffold as no exchange between subunits was observed when preformed coats were incubated with an excess of either SNXs or CSC.. Rigidity of the coat is illustrated by its ability to stabilize membrane tubules pulled out of a GUV. Similar experiments allowed to calculate the polymerization energy of dynamin by plotting membrane tension against the residual force exerted by the membrane tubule carrying the polymerized coat (Roux *et al*, 2010). We measured no significant residual force on SNX/CSC tubules in the range of membrane tension tested.

This suggests a high polymerization energy of the retromer coat which, however, cannot be quantified through this assay at this point. Determining it will require much more work and is beyond the scope of this paper.

In situ, the stability of the coat and its stabilizing effect on the underlying tubule could be relevant in various ways. It might influence endosomal maturation, where efficient intralumenal vesicle formation by the ESCRT complex requires low endosomal membrane tension. Stabilization of membrane tubules through retromer might maintain the endosomal membrane under high tension, which might then suppress MVB formation as long as sufficient amounts of recycling cargo are present in the endosome. Stability of the retromer coat could also be relevant for the final detachment of a coated carrier: It may impose constraints on lipid flow beneath the coat and thereby promote friction-mediated membrane fission (Simunovic *et al*, 2017), when forces pull the tip of the membrane tubule. Such forces might be generated and transmitted through retromer-interacting proteins. In mammalian cells, the CSC interacts with the WASH complex, an activator of Arp2/3 that generates branched actin filaments. It was proposed that actin polymerization generates force to elongate the retromer coated tubule and ultimately cause its fission (Derivery *et al*, 2009; Gomez & Billadeau, 2009; Harbour *et al*, 2012; Phillips-Krawczak *et al*, 2015; Temkin *et al*, 2011; Jia *et al*, 2012). This force would be most efficiently transferred to the growing tubule through a retromer scaffold that is static and stable, which would be in line with our experimental observations. A caveat for this working model is that WASH does not exist in yeast - although also in this organism force produced by actin is harnessed to drive membrane trafficking processes, such as endocytosis (Goode *et al*, 2015; Kaksonen, 2008). One would hence have to postulate that in this system force is transmitted independently of WASH.

The coat radius of 19 nm can be defined by SNX alone, but CSC facilitates membrane constriction at lower SNX concentration. In the constricted zones, the density of occupation of SNX by CSC varies as a function of the starting diameter of the non- constricted membrane tube. This suggests that constriction of less curved membranes engages more CSC, resulting in a SNX coat with a higher degree of CSC-mediated crosslinking between the SNX subunits. The energy provided by these additional bonds and enhanced scaffolding of SNX by CSC may be two factors that enhance the capacity of the coat to work on the membrane. Then, the coat need not operate at a fixed stoichiometry and retromer density but can be tuned according to the circumstances. This could become relevant when coats must be formed on an endosome with higher membrane tension. Tubulation requires more work under these circumstances, which could be provided by forming a coat with higher CSC content. Elevated CSC density should also offer more binding sites for WASH and might thereby enhance force transmission by actin on the growing tubule. Regulating the concentration of active CSC could thereby allow the cell to tune the rate of retromer-mediated membrane exit from endosomes as a function of endosomal membrane tension.

CSC incorporation into the coat may provide additional force for tubule formation. That this contribution depends on CSC oligomerization is underscored by the effect of substitutions in the conserved Vps35-Vps35 interface, which compromise oligomerization (Kendall *et al*, 2020; Leneva *et al*, 2021). They abolish the capacity of CSC to drive membrane constriction, lead to miss-sorting of the retromer-dependent cargo receptor Vps10 in yeast, and, probably therefore, to the observed partial secretion of the vacuolar peptidase CPY (Kendall *et al*, 2020). We did not test the ability of human hVPS35^AAA3KE^ or hVPS35^PISA^ to form a stable coat in vitro, but these variants induced a strong and even dominant negative recycling defect of the retromer cargo GLUT1 in living cells. This points to a conserved role of Vps35-mediated retromer oligomerization in protein recycling from endosomes.

The asymmetry of the Vps35-Vps35 interface was proposed to have potential functional consequences, such as for binding cofactors to the arch in a 1:2 stoichiometry and/or conferring directionality to the growth of the coat (Leneva *et al*, 2021). In our experiments, the coat grew bidirectionally, suggesting that it lacks an inherent preference for adding new subunits at one end. It remains possible, however, that directionality of coat growth can be conferred by additional factors that have not been present in our in vitro system. Structural asymmetry might also be exploited for other purposes, for example for binding cargo such as Vps10, which was proposed to bind to the C-terminal part of Vps35 (Nothwehr *et al*, 1999) and promotes the tubulation activity of SNX/CSC coats (Purushothaman & Ungermann, 2018). Cargo exerting control over tubule formation through CSC recruitment might then ensure that the recycling machinery is activated when needed.

## Material and methods

### Materials

The following lipids were purchased from Avanti Polar Lipids (USA): Egg L-alphaphosphatidylcholine (EPC); 1,2-dioleoyl-sn-glycero-3-phospho-L-serine sodium salt (DOPS); 1,2-dioleoyl-sn-glycero-3-phospho-(1’-myo-inositol-3’-phosphate) (PI3P); 1,2- dioleoyl-sn-glycero-3-phosphoethanolamine-N-(Cyanine 5.5) (Cy5.5 PE); 1-Oleoyl-2-[12- [(7-nitro-2-1,3-benzoxadiazol-4-yl)amino]dodecanoyl]-sn-Glycero-3-Phosphocholine (NBDPC). All lipids were dissolved in chloroform. Phosphatidylinositol phosphates were dissolved in chloroform/methanol/water (20:10:1). Texas red DHPE (Thermofisher cat. T1395MP) was purchased as a mixed isomer. The para isomer was separated by thin layer chromatography as previously described (Dar *et al*, 2015).

### Cell culture, strains and plasmids

Yeast cells BY4742 were grown at 30°C in YPD (2% peptone, 1% yeast extract, 2% ý-D- glucose). Genes were deleted by replacing a complete open reading frame with a marker cassette (Janke *et al*, 2004; Güldener *et al*, 1996) (see Appendix Table S1 for a list of strains used in this study and Appendix Table S2 for a list of PCR primers used in this study). Gene tagging was done as described (Sheff & Thorn, 2004). Strains used for expression and purification of the retromer complex have been previously described (Appendix Table 1). VPS35 with genomic mutations at the C-terminus were amplified by PCR from a synthetic gene corresponding to the last 1000 bp of VPS35 for the PISA mutant, or from the pRS315-Vps35AAA3KE plasmid (Kendall *et al*, 2020) for the AAA3KE mutant. These fragments were then fused to a LEU2 cassette by fusion PCR and transformed into yeast cells (Janke *et al*, 2004)

### Live microscopy

Vacuoles were stained with FM4-64 essentially as described (Desfougères *et al*, 2016). An overnight preculture in HC (Hartwell’s complete) medium was used to inoculate a 10 ml culture. Cells were then grown in HC to an OD_600_ between 0.6 and 1.0. The culture was diluted to an OD_600_ of 0.4, and FM4-64 was added to a final concentration of 10 µM from a 10 mM stock in DMSO. Cells were labelled for 60 min with FM4-64, washed three times in fresh media by short and gentle centrifugation in a benchtop centrifuge, and then incubated for 60 min in media without FM4-64. Right before imaging, cells were concentrated by a brief low-speed centrifugation, resuspended in 1/10 of their supernatant, placed on a glass microscopy slide and overlaid with a 0.17 mm glass coverslip. Imaging was done with a NIKON Ti2 spinning disc confocal microscope with a 100x 1.49 NA lens.

Z-stacks were taken with a spacing of 0.3 µm and assembled into maximum projections. Image analysis was performed with ImageJ. Pearson’s correlation coefficient was used to quantify the colocalization between Vps10 and FM4-64. The Nikon NIS-Elements Software Pearson’s correlation tool was used on at least 5 single stacks containing at least 100 cells each. All performed experiments were repeated at least three times. SEM calculation and potting were done with Graphpad PRISM software.

### Protein purification

TAP-tagged retromer complex was extracted from yeast as previously described (Purushothaman *et al*, 2017; Purushothaman & Ungermann, 2018). Briefly, a 50 mL preculture of cells was grown over night to saturation in YPGal medium. The next day, two 1L cultures in YPGal were inoculated with 15 mL of preculture and grown for 20 h to late log phase (OD_600_ = 2 to 3). All following steps were performed at 4°C. Cells were pelleted and washed with 1 pellet volume of cold RP buffer (retromer purification buffer: 50 mM Tris pH 8.0, 300 mM NaCl, 1 mM MgCl_2_, 1 mM PMSF, Roche complete protease inhibitor).

Pellets were either processed immediately or flash-frozen in liquid nitrogen and stored at - 80°C. For cell lysis, the pellet was resuspended in one volume of RP buffer and passed through a French press (One shot cell disruptor, Constant Systems LTD, Daventry, UK) at 2.2 Kpsi. 1 mg DNAse I was added to the lysate followed by a 20 min incubation on a rotating wheel. The lysate was precleared by centrifugation for 30 min at 45’000 xg in a Beckman JLA 25.50 rotor and cleared by a 60 min centrifugation at 150’000 xg in a Beckman Ti60 rotor. The cleared supernatant was passed through a 0.2 µm filter and transferred to a 50 mL Falcon tube. 1 mL IgG bead suspension (GE Healthcare, cat 17- 0969-01) was washed three times with RP buffer and added to the supernatant. After 60 min incubation on a rotating wheel, beads were spun down and washed 3 times with RP buffer. 250 µg of purified HIS-TEV protease from E. coli was added to the beads. After 30 min incubation at 4°C, beads were centrifuged, the supernatant containing purified retromer subcomplex was collected and concentrated on a 100 kDa cutoff column (Pierce™ Protein Concentrator PES, 100K MWCO). The concentrated protein fraction was re-diluted in RP buffer and reconcentrated 3 times. This final step allowed for removal of TEV protease and a high enrichment for intact complexes. Proteins were concentrated to ∼2 mg/mL, aliquoted in 10 µL fractions and flash-frozen in liquid nitrogen. Proteins were stored at -80°C and used within 3 months. Thawed aliquots were used only once.

### Supported membrane tubes

Supported membrane tubes were generated as described (Dar *et al*, 2015).

Briefly, glass coverslips were first washed with 3 M NaOH for 5 min and rinsed with water before a 60 min treatment with piranha solution (95% H2SO4 / 30% H2O2 3:2 v/v).

Coverslips were rinsed with water and dried on a heat block at 90°C. Coverslips were then silanized with 3-glycidyloxypropyltrimethoxysilane (Catalogue no. 440167, Sigma) for 5 h under vacuum, rinsed with acetone and dried. Polyethylene glycol coating was done by placing the coverslips in a beaker containing PEG400 (Sigma) at 90°C for 60 h. Coverslips were washed with distilled water and stored for up to 2 months at room temperature in a closed container.

To generate supported membrane tubes, lipids were mixed from 10 mg/mL stocks in a glass vial and diluted to a final concentration of 1 mg/mL in chloroform. The same lipid mix was used throughout this study (5% PI(3)P, 15% DOPS, 0.1% fluorescent lipid tracer (in most cases Texas red DHPE), 79.5% egg-PC). Lipids were then spotted (typically 1 µL, corresponding to about 1 nmol) on the coverslips and dried for 30 min under vacuum. The coverslip was mounted on an IBIDI 6-channel µ-slide (µ-Slide VI 0.4. IBIDI, catalog no: 80606). Lipids were hydrated for 15 min with buffer (PBS) and SMTs were generated by injecting PBS into the chamber using an Aladdin Single-Syringe Pump (World Precision Instruments, model n°. AL-1000) at a flow rate of 1,5 mL/min for 5 min. SMTs were left to stabilize without flow for 5 min before the start of the experiment. Protein stocks (typically 1-2 µM) were first diluted in PBS and then injected in the chamber at a flow rate of 80 µL per minute. Tubes were imaged with a NIKON Ti2 spinning disc confocal microscope equipped with a 100x 1.49 NA objective.

### Native-PAGE, SDS-PAGE and Western blotting

For analysis of CSC oligomer formation, 10 μL of purified CSC (∼2 mg/mL) were diluted 1:1 with water and incubated for 5 min at 25°C. Samples were run on a commercial native- PAGE gel (3 to 12%, Bis-Tris, 1.0 mm, Mini Protein Gel, 10-well, Invitrogen, Cat. BN1001BOX) at 100 V tension using as running buffer 50 mM BisTris, 50mM Tricine, pH 6.8 (Invitrogen number BN2007). Cathode buffer: running buffer + 1/200 0.4% Coomassie G-250. Sample buffer: 50 mM BisTris, 6 N HCl, 50 mM NaCl, 10% w/v Glycerol, 0.001% Ponceau S, pH 7.2). Gels were run at 4° C. After the run, gels were then washed in 20% ethanol + 10% acetic acid for 2 hours.

### Quantifcation of SMT fluorescence

SMT fluorescence was quantified with ImageJ. Line scan analysis was performed along tubules using an ImageJ plugin (available as a txt file in the supplement; see also Suppl. Fig.2). Each line scan was performed perpendicular to the tubule. Scans were made along the tubule with a one-pixel increment. For each line scan, a Gaussian curve was fitted, and the maximum height was extracted. Maximum height was then plotted against the tube length for all channels. For quantification of the diameters of the tubes, lipid fluorescence values of a tubule underneath a constricted protein domain, extracted from the series of line scans described above, was sorted in ascending order. The curve typically showed two plateaus, the lower corresponding to the constricted state and the higher to the non- constricted one. Plotting the corresponding GFP values confirmed that the GFP-labelled protein localized to the constricted zone. For each tube, the zones corresponding to the constricted and non-constricted areas were determined manually and the mean fluorescence value was used to calculate the tube diameter. Tube diameter was calibrated as described (Dar *et al*, 2015) using purified human ΔPRD-dynamin-1 (Colom *et al*, 2017) as a reference.

### Tube Pulling

The experimental set-up used to aspirate GUVs with a micropipette and pull a membrane tube was the same as previously reported (Chiaruttini *et al*, 2015) combines bright-field imaging, spinning disc confocal microscopy and optical tweezers on an inverted Nikon Eclipse Ti microscope. GUVs were made by electro-formation as described (Angelova *et al*, 1992). Briefly, lipid mix (the same mix as for SMT experiments, supplemented with 0.03% mol/mol of the biotinylated lipid DSPE-PEG2000-Biotin, Avanti Polar Lipids, Alabaster, AL, USA) in chloroform was deposited on indium-titanium oxide glass slides and dried for 60 min at 55°C to evaporate all solvents. GUVs were electroformed at 1 V and 10 Hz for 60 min at 55°C in a 380 mM sucrose solution. GUVs were then removed from the chamber and placed in an Eppendorf tube until use. GUVs were used within 1–2 h after formation. A GUV is aspirated within a micropipette connected to a motorized micromanipulator (MP-285, Sutter Instrument, Novato, CA, USA) and a homemade pressure control system (Zaber Micro linear actuator, Zaber Technologies Inc., Canada) that sets the aspiration pressure ΔP. Then, a membrane nanotube is pulled out from the vesicle through a streptavidin-coated bead (3.05 μm diameter, Spherotec, Lake Forest, IL, USA) held in a fixed optical trap. The optical trap was custom-made with a continuous 5 W 1064 nm fiber laser (ML5-CW-P-TKS-OTS, Manlight, Lannion, France) focused through a 100X 1.3 NA oil immersion objective. The force F exerted on the bead was calculated from Hooke’s law: F = k*Δx, where k is the stiffness of the trap (k = 60 pN.μm^−1^) and Δx the displacement of the bead from its equilibrium position. A mix of SNXs / CSC-mClover at 1 µM with 280 mosm osmolarity was injected with a micropipette connected to a motorized micromanipulator and to the Fluigent pressure control system (MFCS-VAC, −69 mbar; Fluigent).

### CLEM

CLEM was performed as described (Muriel *et al*, 2021; Kukulski *et al*, 2012). Briefly, cells of a logarithmically growing culture were concentrated by centrifugation at 3000 rpm for 2 min at RT. A few microliters of a thick cell slurry were pipetted onto a 3-mm-wide and 0.1- mm deep specimen carrier (Wohlwend type A) closed with a flat lid (Wohlwend type B).

The assembled carrier sandwich was high-pressure frozen using a Wohlwend HPF Compact 02 and disassembled in liquid nitrogen. High-pressure frozen samples were processed by freeze substitution and embedding in Lowicryl HM20 using the Leica AFS 2 robot as described (Kukulski *et al*, 2012). 300 nm sections were cut with a diamond knife using a Leica ultramicrotome, collected in water and picked up on carbon-coated 200- mesh copper grids (AGS160; Agar Scientific). For light microscopy the grid was placed onto a drop of water and mounted onto a microscopy slide. Light microscopy images were acquired on a NIKON Ti2 spinning disc confocal microscope with a 100x 1.49 NA lens. The grid was recovered, dried and stained with Reynolds lead citrate for 10 min. 10-nm protein A-coupled gold beads were adsorbed to the top of the section as fiducials for tomography. TEMs were acquired on a FEI Tecnai 12 at 120 kV using a bottom mount FEI Eagle camera (4k x 4k). For tomographic reconstruction, tilt series were acquired over a tilt range of ± 60° at 1° increments using the Serial EM software. Tomogram reconstruction was performed using the IMOD software package with gold fiducial alignment.

### Mammalian cell experiments

All chemical reagents were from Sigma-Aldrich unless specified otherwise. Other reagents: Opti-MEM (Thermo Fischer, 11058021) and Trypsin (Thermo Fischer, 27250018); LysoTracker® Deep Red (Thermo Fisher Scientific, L12492; Protease inhibitor (PI) cocktail (final concentrations: 40 M pefablock SC (Merck, 11429876001), 2.1 M leupeptin (Merck, 11529048001), 80 μM o-phenantroline (Merck, 131377), 1.5 μM pepstatin A (Merck, 11524488001).

#### Cell culture, transfection and treatments

HK2 cells were grown in DMEM-HAM’s F12 (GIBCO-Life Technologies); supplemented with 5% fetal calf serum, 50 IU/mL penicillin, 50 mg/mL streptomycin, 5 μg/mL insulin, 5 μg/mL transferrin, 5 ng/mL selenium (LuBio Science). Cells were grown at 37°C in 5% CO2 and at 98% humidity. Media, serum and reagents for tissue culture were purchased from GIBCO (Invitrogen). HK2 cells were transfected with different plasmids using X- tremeGENE HP DNA transfection reagent (Sigma-Aldrich) according to the manufacturer’s instructions. Briefly, the plasmid was diluted with Opti-MEM I Medium without serum to a final concentration of 1 μg plasmid DNA /100 μl medium (0.01 μg/μl) and gently mixed.

Then, 3 μl of X-tremeGENE HP DNA Transfection Reagent was added directly into the medium containing the diluted DNA. The transfection reagent: DNA complex was incubated for 30 at room temperature under the hood. Finally, the transfection complex was added to the cells in a dropwise manner and they were incubated 24 hours at 37°C in a CO_2_ incubator.

The HK-2 cell line was checked for mycoplasma contamination by a PCR-based method. All cell-based experiments were repeated at least three times.

#### Knockouts and RNA interference

For RNA interference, HK2 cells were plated in 24well-plate and then transfected with siRNA using Lipofectamine RNAiMax (Thermo Fisher Scientific). For each well to be transfected, was first prepared the RNAi duplex-Lipofectamine RNAiMAX complexes as follows: 6 pmol of RNAi duplex were diluted in 100 µl Opti-MEM I Medium without serum in the well of the culture plate and gently mixed. Then, 1 µl Lipofectamine RNAiMAX was added to each well containing the diluted RNAi molecules, gently mixed and incubated for 20 minutes at room temperature under sterile conditions. In that time cells were detached, counted, and diluted in complete growth medium without antibiotics so that 500 µl contains the appropriate number of cells to give 30% confluence 24 hours after plating. After the 20 minutes of incubation at room temperature to each well with RNAi duplex - Lipofectamine RNAiMAX complexes were added 500 µl of the diluted cells. This gives a final volume of 600 µl and a final RNA concentration of 10 nM. The 24well-plate was gently mixed gently by rocking and incubated 24-72 hours at 37°C in a CO2 incubator.

The siRNA targeting VPS35 was from Sigma (5’ CTGGACATATTTATCAATATA 3’; 3’ TA- TATTGATAAATATGTCCAG 5’). It was used at 10 nM final concentration. Control cells were treated with identical concentrations of siGENOME Control Pool Non-Targeting from Dharmacon (D-001206-13-05).

#### Immunostaining

HK2 cells were grown to 70% confluence on glass coverslips before immunofluorescence microscopy was performed. Cells were fixed for 10 min in 4% paraformaldehyde in PBS (phosphate-buffered saline). After fixation, cells were incubated (30 min at room temperature) in blocking buffer with (permeabilized cells) or without (non-permeabilized cells) 0.05% (w:v) saponin (Sigma-Aldrich, 558255), 0.5% (w:v) BSA and 50 mM NH4Cl in PBS. The cells were incubated for 1 h with primary antibody in blocking buffer, washed three times in PB and incubated for 1 h with the secondary antibody in blocking buffer. Then, cells were washed three times in PBS, mounted with Mowiol (Sigma-Aldrich, 475904-M) on slides and analysed by confocal microscopy.

Primary antibodies were anti-LAMP1 (H4A3, USBiologicvak Life Sciences) and anti-Glut1 (ab15309 Abcam). Secondary antibodies were Cy3-conjugated AffiniPure Donkey antiMouse IgG H+L (Jackson Immuno Research); Cy3-conjugated AffiniPure Donkey antiRabbit IgG H+L (Jackson Immuno Research); Alexa fluor®488-conjugated AffiniPure Donkey anti-Rabbit IgG H+L (Jackson Immuno Research).

*Confocal fluorescence microscopy and image processing*.

Confocal microscopy was performed on an inverted confocal laser microscope (Zeiss LSM 880 with airyscan) with a 63x 1.4 NA oil immersion lens. Z-stack Images were acquired on a Zeiss LSM880 microscope with Airyscan. GLUT1-fluorescence was quantified using ImageJ. Individual cells were selected using the freeform drawing tool to create a ROI (ROI). The ‘Measure’ function provided the area, the mean grey value and integrated intensity of the ROI. The mean background level was obtained by measuring the intensity in three different regions outside the cells, dividing them by the area of the regions measured, and averaging the values obtained. This background noise was removed from each cell, yielding the CTCF (corrected total cell fluorescence): CTCF=integrated intensity of cell ROI − (area of ROI × mean fluorescence of background).

To quantify the degree of co-localisation, confocal z-stacks were acquired. Single channels from each image in 8-bit format were thresholded to subtract background and then the “Just Another Colocalisation Plug-in” (JACOP) of ImageJ was used to measure the Pearson’s correlation coefficient.

### Gel electrophoresis and Western blot

Ctrl and Vps35-KD HK2 cells were plated into 12-well tissue culture test plates (TPP) until 72h after transfection with the siRNAs. Cells were then washed three times with ice-cold PBS, scraped, and proteins were extracted in ice-cold lysis buffer (150 mM NaCl, 2 mM EDTA, 40 mM HEPES, and 1% Triton X-100) supplemented with phosphatase (Roche #04906837001) and protease inhibitor cocktail. Protease inhibitor (PI) cocktail (final concentrations: 40 μM pefablock SC (Merck, 11,429,876,001), 2.1 μM leupeptin (Merck, 11,529,048,001), 80 μM o-phenantroline (Merck,131,377), 1.5 μM pepstatin A (Merck, 11,524,488,001). Protein extracts were supplemented with 1/4 volume of 5x reducing sample buffer (250 mM Tris-Cl, pH 6.8, 5% β-mercaptoethanol, 10% SDS, 30% glycerol, 0.02% bromophenol blue) and heated to 95 °C for 5 min. The samples were run on SDSpolyacrylamide gels (W x L x H: 8.6 x 6.8 x 0.15 cm). Running gels were either 8% or 416% protogel (30% w/v acrylamide, 0.8% bisacrylamide (37.5:1 solution, National diagnostics, Atlanta, USA),, 0.38 M Tris, pH 8.8, 0.1% w/v SDS (Applichem, 475904-M), 0.06% TEMED (Applichem, A1148), 0.06% w/v APS (Applichem, A2941). The stacking gels were prepared as follows: 6% acrylamide, 0.16% bis-acrylamide, 0.1 M Tris, pH 6.8, 0.1% SDS, 0.1% TEMED, 0.05% ammonium persulfate. The gels were run at constant current (35 mA). Proteins were blotted onto 0.45 µm nitrocellulose membrane (Amersham) overnight at a constant current of 200 mA using a Trans-Blot® Cell (Bio-Rad, USA).

After incubation with the primary antibody, signals were detected by secondary antibodies coupled to infrared dyes (LI-COR) and detected on a LI-COR Odyssey Infrared Imager.

Images were exported as TIFF files and processed in Adobe Photoshop. Band intensity was quantified using ImageJ band analysis (Schneider CA et al., 2012). We used antiLAMP1 (H4A3, USBiologicvak Life Sciences), anti-Tubulin (T9026 Sigma-Aldrich) antiVps35 (ab10099 Abcam, ab157220 Abcam).

### Statistics

Where averages were calculated, the values stem from experiments that were performed independently. For all experiments, significance of differences was tested by a two-tailed ttest.

## Acknowledgements

We thank Christian Ungermann for strains expressing CSC and SNX. This work was supported by grants from the SNSF (179306 and 204713) and ERC (788442) to AM.

## Supplementary Figures

**Suppl. Figure 1:**
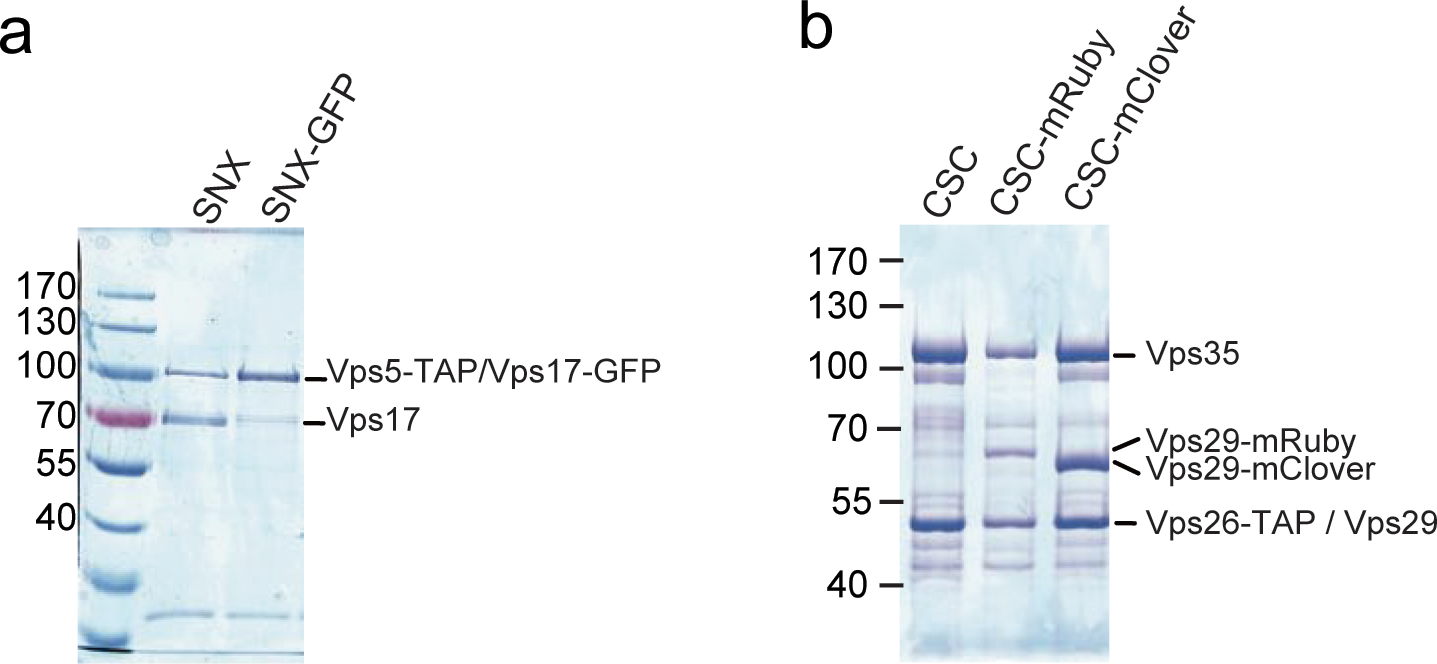
SNX and CSC proteins extracted from yeast. The indicated TAP-tagged protein complexes were extracted from yeast and analyzed by SDS-PAGE and Coomassie staining.

**Suppl. Figure 2:**
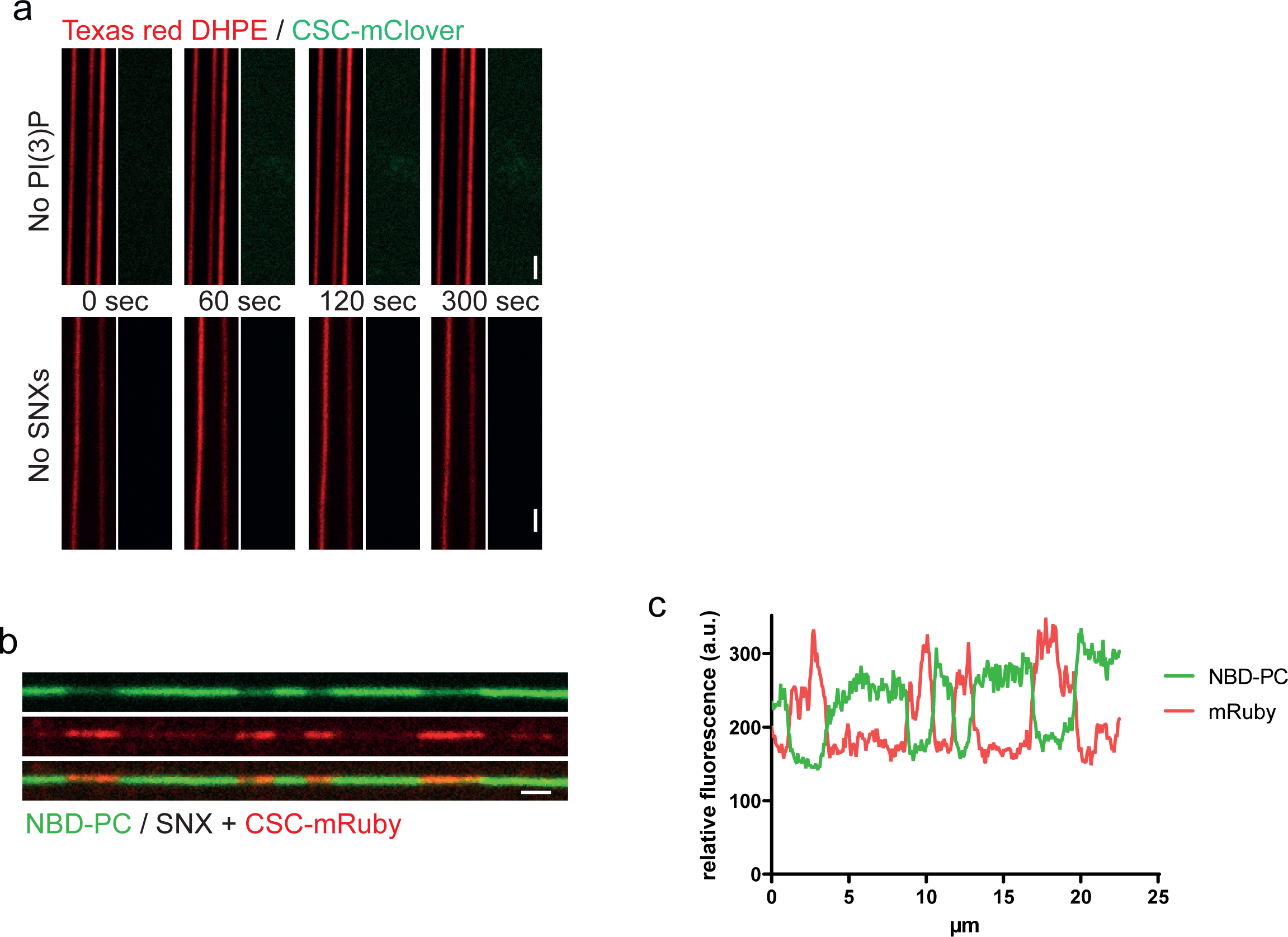
Lipid and SNX dependence of CSC binding and constriction of tubules. **a.** PI3P and SNX dependence of CSC recruitment. SMTs were prepared with or without PI3P, incubated with 25 nM CSC^mClover^ in the presence or absence of 25 nM SNX complex and analyzed by fluorescence microscopy. Scale bars: 2 µm. **b.** Coat formation and tube constriction shown through NBD-PC, a lipid labelled at a fatty acyl chain. SMTs labelled with NBD-PC instead of Cy5.5-PE were incubated with SNX and CSC^mClover^ as in Fig. 1a and analyzed by fluorescence microscopy. **c.** Line scan analysis of the tubule shown in b.

**Suppl. Figure 3:**
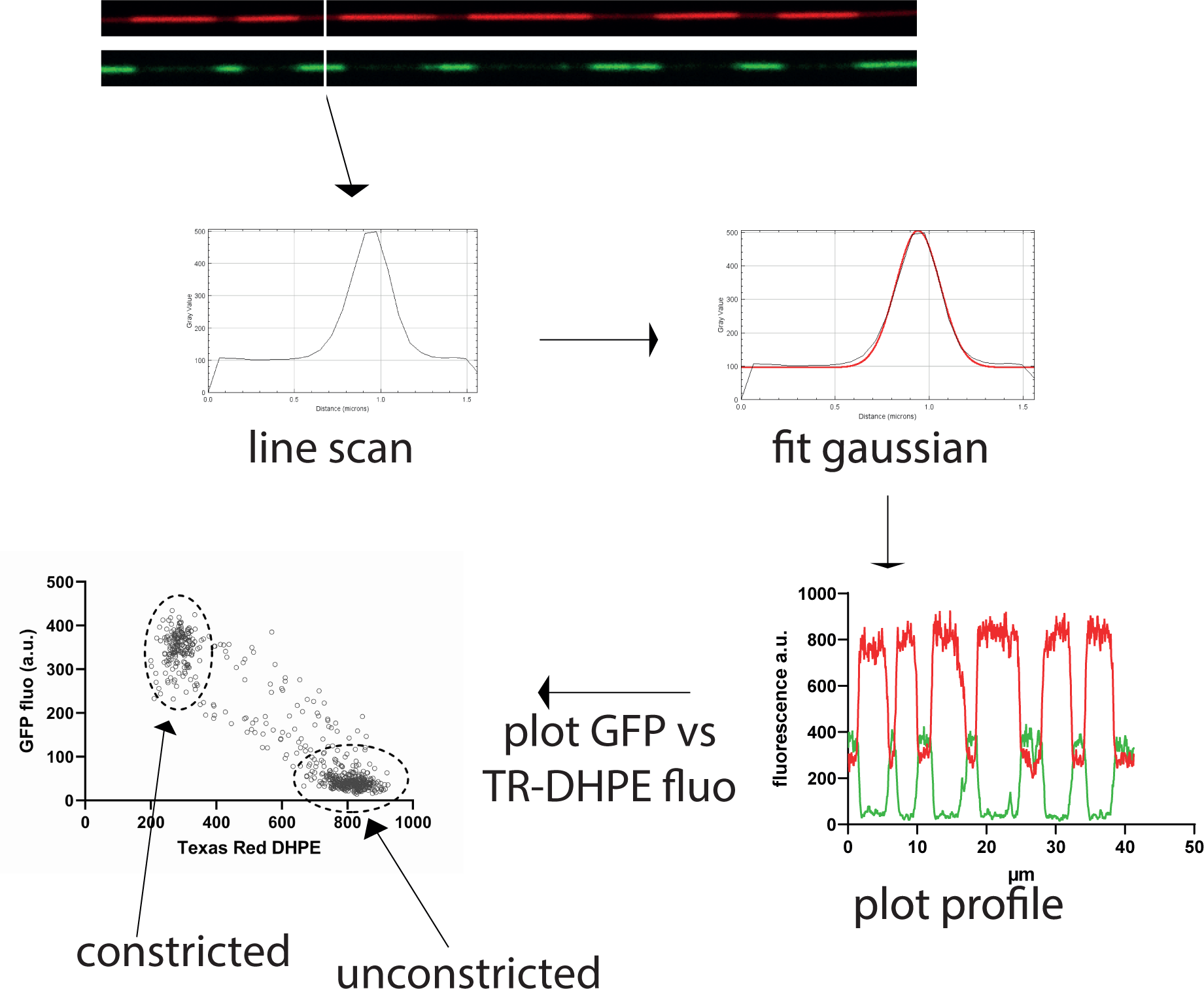
Procedure for quantifying constrictions along tubules. A series of line scans were performed along the entire length of an SMT, perpendicular to the tube. For each line scan, a Gaussian curve was fitted, and the maximum height of the coat signal (e.g. GFP) and the lipid signal (e.g. Texas Red DHPE) were extracted. The maximum height was then plotted against the intensity of the lipid signal along the same scan. This was performed for every line scan along the tube, yielding two populations. The number of dots in each population are a measure for the length of constricted and non- constricted regions. For each tube, the zones corresponding to the constricted and non- constricted areas were determined manually and their mean fluorescence value was used to calculate the tube diameter by comparison with the signals obtained with dynamincoated tubes.

**Suppl. Figure 4:**
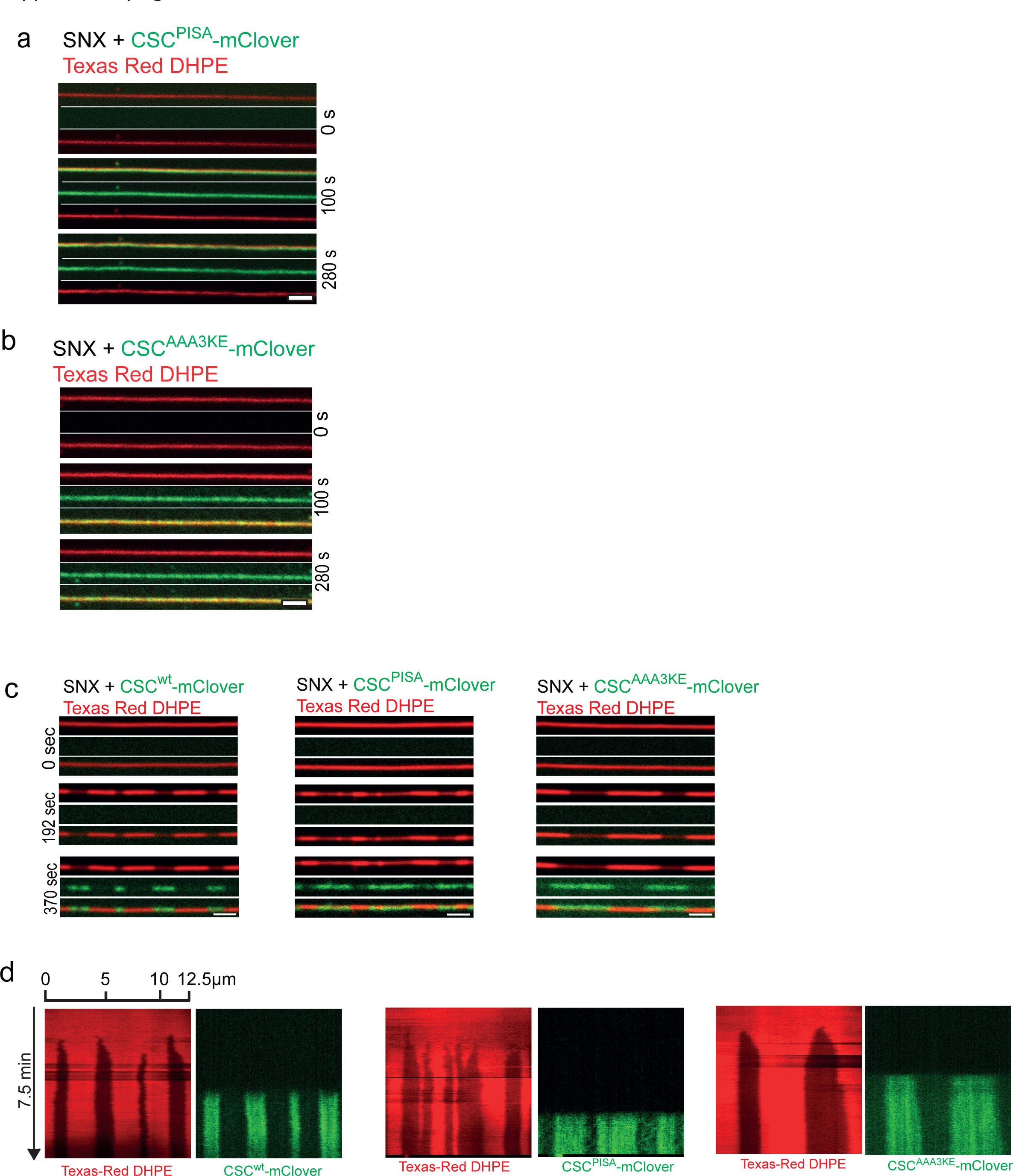
Lack of constricted domain formation after co-incubating SNX with CSC containing *vps35^PISA^* or *vps35^AAA3KE^*. **a, b.** SMT assay with the dimerization mutants. 50 nM SNX and 50 nM of CSC^mClover^ carrying the indicated Vps35 variants were incubated with SMTs and imaged by confocal microscopy. Scale bars: 2 µm **c**. Binding of CSC variants to preformed SNX domains. SMTs were incubated with high SNX concentration (100 nM) for 2 min to pre-form constrictions zones. Unbound SNX was washed away and 50 nM CSC^mClover^ containing the indicated Vps35 variants was added. Panels show tubes before SNX addition (0 sec), after the formation of SNX only coats (192 sec) and after incubation with CSC^mClover^ variants (370 sec). Tubes were imaged by spinning disc confocal microscopy at 0.5 Hz throughout the experiment. Scale bars: 2 µm **d.** Kymographs of the tubes shown in c.

**Suppl. Figure 5.**
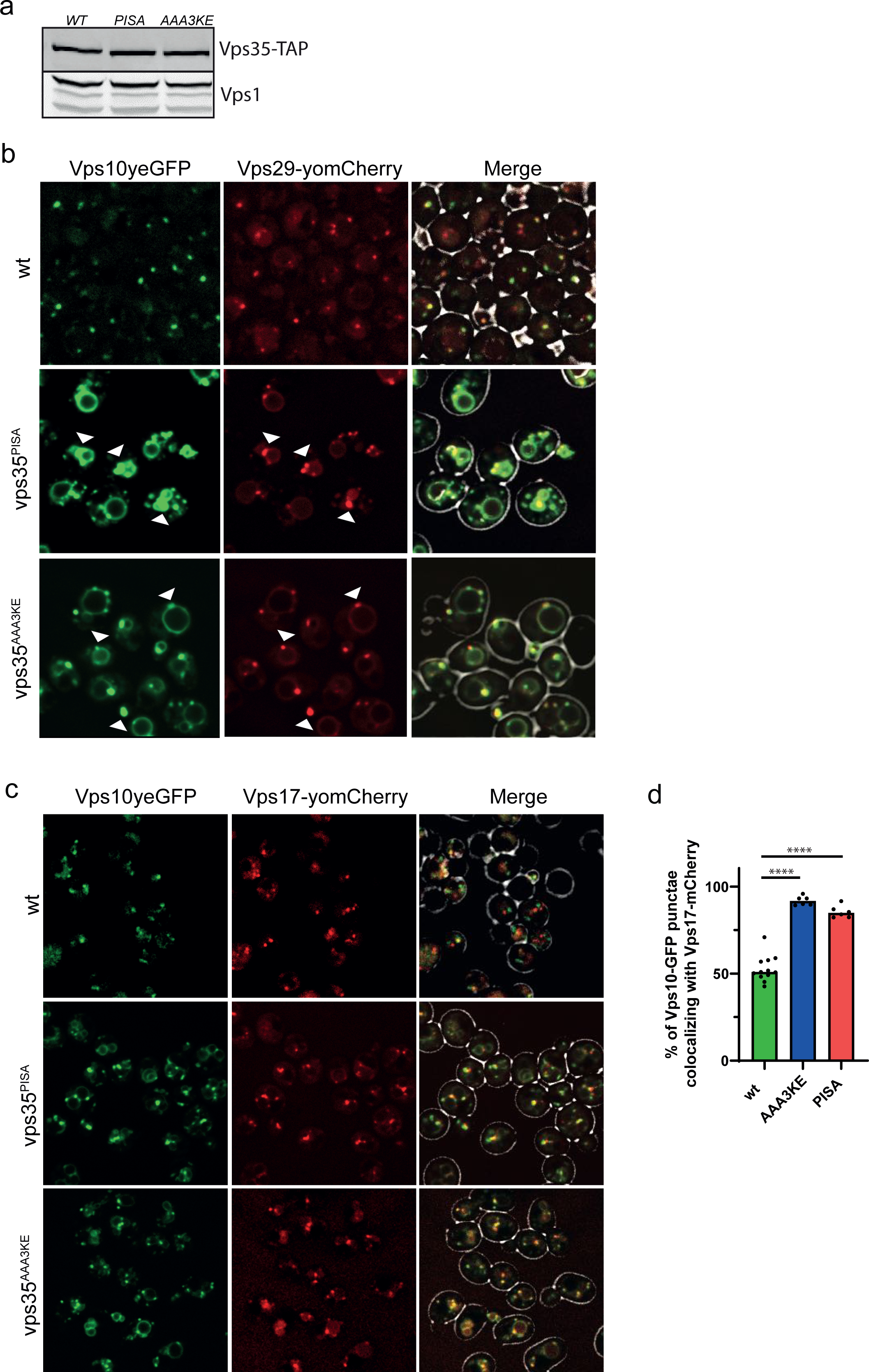
Effect of vps35 dimerization mutants on the localization of Vps10, Vps17 and Vps29 in yeast cells. **a**. Vps35 expression levels. Cells expressing the indicated VPS35 alleles as TAP fusion proteins were grown in YPD to mid-logarithmic phase. Whole-cell extracts were analyzed by SDS-PAGE and Western blotting against the TAP tag. **b, c.** Colocalization of Vps10^yEGFP^ with yomCherry fusions of (b) Vps29 and (c) Vps17. All tags were introduced at the genomic locus, generating C-terminal fusions as the sole source of the respective proteins. Cells had been harvested in logarithmic growth (OD600nm between 0.8 and 1.0), concentrated in their medium by a short spin and immediately imaged by spinning disc confocal microscopy. Arrowheads indicate co-localization of retromer subunits with Vps10^yEGFP^. The merge includes a brightfield image to indicate the cell boundaries. **d.** Quantification of Vps10^yEGFP^ dots colocalizing with Vps17^mCherry^ in cells from c. Quantification was done with the Nikon analysis software NIS-Elements. Statistical significance was assessed with a two-tailed t-test. **** P < 0.0001.

**Suppl. Figure 6.**
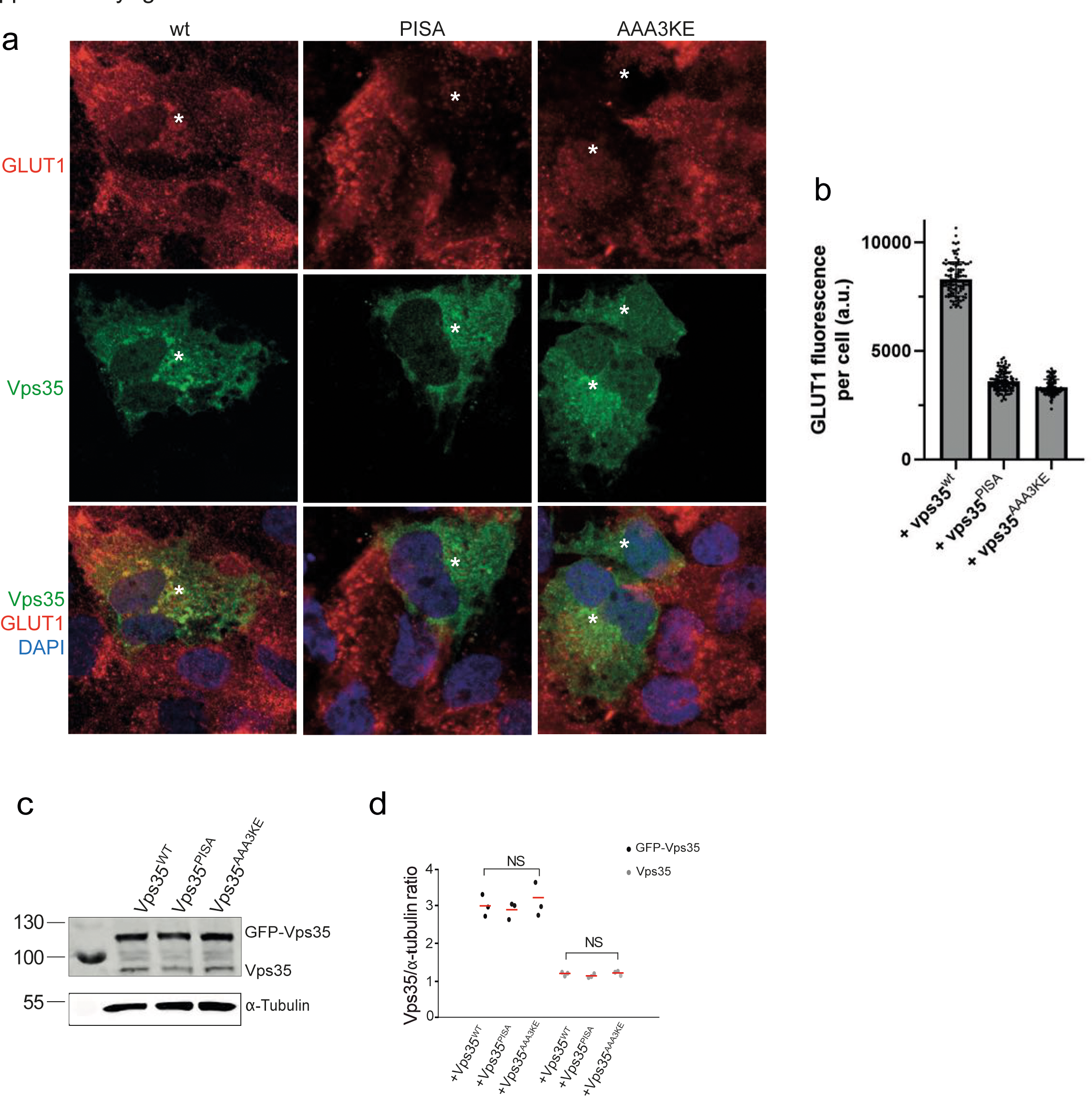
Dominant negative effect of hVPS35 variants on GLUT1 sorting in HK2 cells. **a.** HK2 cells were transfected with a plasmid carrying the indicated variants of hVPS35^GFP^, or with an empty plasmid, and cultivated for 24h. They were fixed, stained with antibody to GLUT1 (red) and with DAPI (blue) and analyzed by confocal microscopy. The cells were not detergent permeabilized. Scale bar: 10 µm **b.** Quantification of GLUT1 fluorescence in the cells from a was performed as in Fig. 9c. **c.** Expression of Vps35 variants in cells from a was analyzed by SDS-PAGE and Western blot against Vps35. Tubulin served as loading control. **d.** Quantification of the Vps35/tubulin ratio in cells from a. Data stems from three independent experiments and show the mean. NS: not significant according to a two tailed ttest (p>0.01).

**Suppl. Figure 7:**
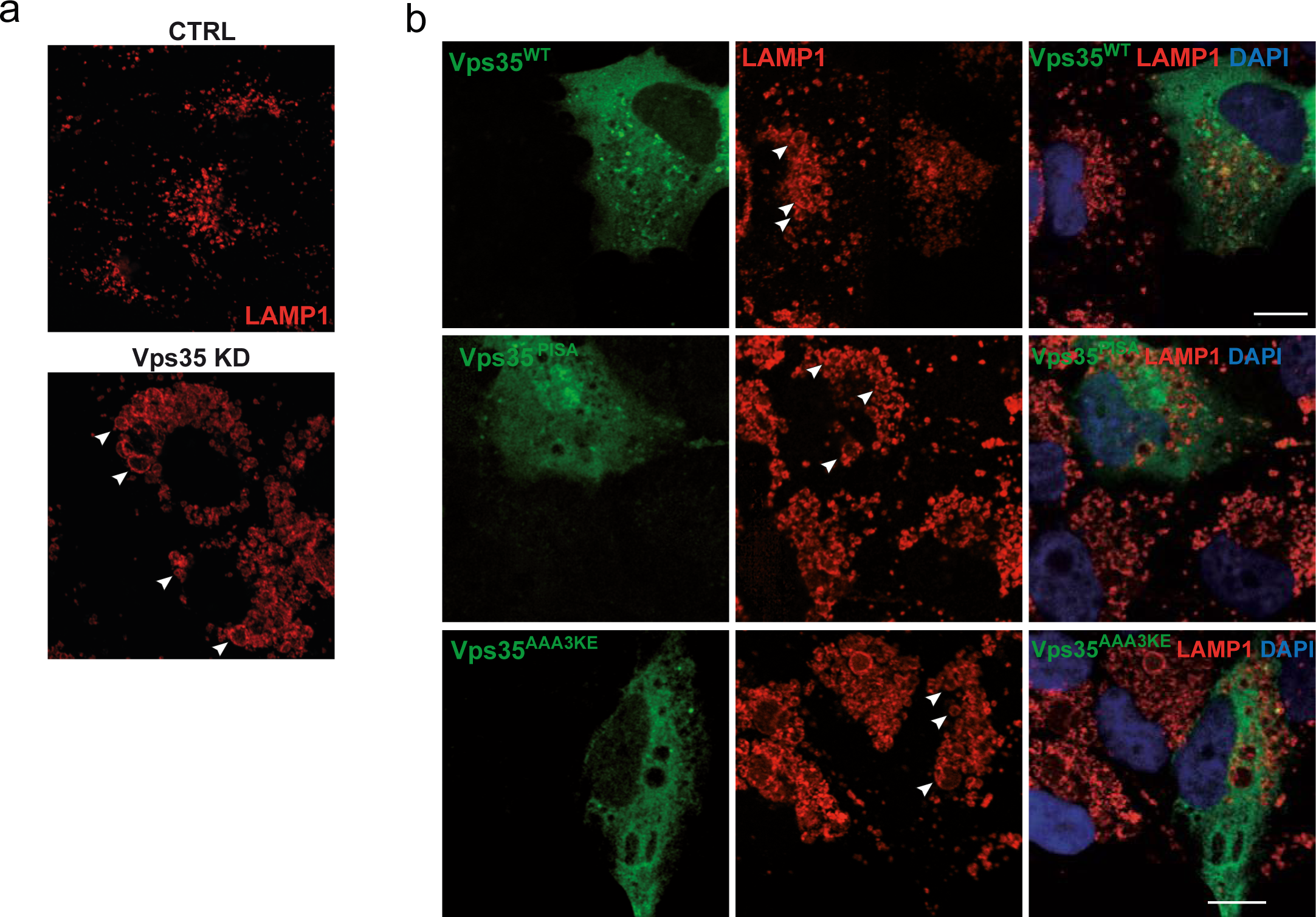
Effect of VPS35 variants on the size of LAMP1 compartments. **a.** Effect of VPS35 knockdown. HK2 cells were treated with siRNA targeting VPS35 or with mock siRNA. The cells were fixed and immuno-stained with anti-LAMP1 antibody and DAPI and imaged by confocal microscopy. Representative images are shown. Scale bar: 10 µm. Arrowheads point to examples of enlarged LAMP1-compartments. **b.** Effect of VPS35 variants. Vps35 knock-down cells were transfected with a plasmid expressing siRNA-resistant wildtype or mutant forms of VPS35^GFP^, or with an empty plasmid. Cells were fixed, immuno-stained with anti-LAMP1 and DAPI and imaged by confocal microscopy. Scale bar: 10 µm. Arrowheads point to examples of enlarged LAMP1-compartments.

## Supplementary Tables

**Suppl. Table 1.** strains used in this study.

| Strain | Genotype | Reference |
| --- | --- | --- |
| CUY105 | *MATa his3Δ200 leu2Δ0 met15Δ0 trp1Δ63 ura3Δ0* | (Purushothaman & Ungermann, 2018; Purushothaman *et al*, 2017) |
| CUY100 | *MATalpha his3Δ200 leu2Δ0 lys2Δ0 met15Δ0 trp1Δ63 ura3Δ0* | (Purushothaman & Ungermann, 2018; Purushothaman *et al*, 2017) |
| CUY9228 | CUY105, *VPS5pr::natNT1-GAL1pr VPS17pr::kanMX-GAL1pr VPS5::TAP-URA3 vps35::HIS3* | (Purushothaman & Ungermann, 2018; Purushothaman *et al*, 2017) |
| CUY9495 | CUY100, *VPS26pr::HIS3-GAL1pr VPS29pr::natNT2-GAL1pr VPS35pr::hphNT1-GAL1 VPS35::TAP-kanMX vps5::TRP1 vps17::LEU2* | (Purushothaman & Ungermann, 2018; Purushothaman *et al*, 2017) |
| CUY9711 | CUY105, *VPS5pr::natNT1-GAL1pr VPS17pr::kanMX-GAL1pr VPS5::TAP-URA3 vps35::HIS3 VPS17::GFP-hphNT1* | (Purushothaman & Ungermann, 2018; Purushothaman *et al*, 2017) |
| CUY9932 | CUY105, *VPS26pr::HIS3-GAL1pr VPS29pr::natNT2-GAL1pr VPS35pr::hphNT1-GAL1 VPS29-mClover::kan VPS26::TAP-URA3 vps5Δ::TRP1* | (Purushothaman & Ungermann, 2018; Purushothaman *et al*, 2017) |
| CUY9935 | CUY105, *VPS26pr::HIS3-GAL1pr VPS29pr::natNT2-GAL1pr VPS35pr::hphNT1-GAL1pr VPS29-mCherry::kan VPS26::TAP-URA3 vps5Δ::TRP1* | (Purushothaman & Ungermann, 2018; Purushothaman *et al*, 2017) |
| NG403 | CUY9932, Vps35 E681A/E682A/D686A/K726E/K729E/K730E (AAA3KE) | This study |
| NG406 | CUY9932, Vps35 D671A/L772A/K726A/R775A (PISA) | This study |
| 4510 | BY4741 WT *vps10yeGFP::His* | This study |
| 4511 | *BY4741 vps35Δ::KanMX Vps10yeGFP::spHIS5* | This study |
| 4512 | *BY4741 VPS35PISA::LEU Vps10yeGFP::spHIS5* | This study |
| 4513 | *BY4741 VPS35AA3KE::LEU Vps10yeGFP::spHIS5* | This study |
| 4643 | *BY4741 WT VPS10yeGFP::spHIS5 VPS17yomCherry::CaURA3* | This study |
| 4645 | *BY4741 VPS35PISA::LEU VPS10yeGFP::spHIS5*<br>*VPS17yomCherry::CaURA3* | This study |
| 4646 | *BY4741 VPS35AA3KE::LEU VPS10yeGFP::spHIS5*<br>*VPS17yomCherry::CaURA3* | This study |
| 4647 | *BY4741 WT VPS10yeGFP::spHIS5 VPS29yomCherry::CaURA3* | This study |
| 4649 | *BY4741 VPS35PISA::LEU VPS10yeGFP::spHIS5*<br>*VPS29yomCherry::CaURA3* | This study |
| 4650 | *BY4741 VPS35AA3KE::LEU VPS10yeGFP::spHIS5*<br>*VPS29yomCherry::CaURA3* | This study |

**Suppl. Table 2.**
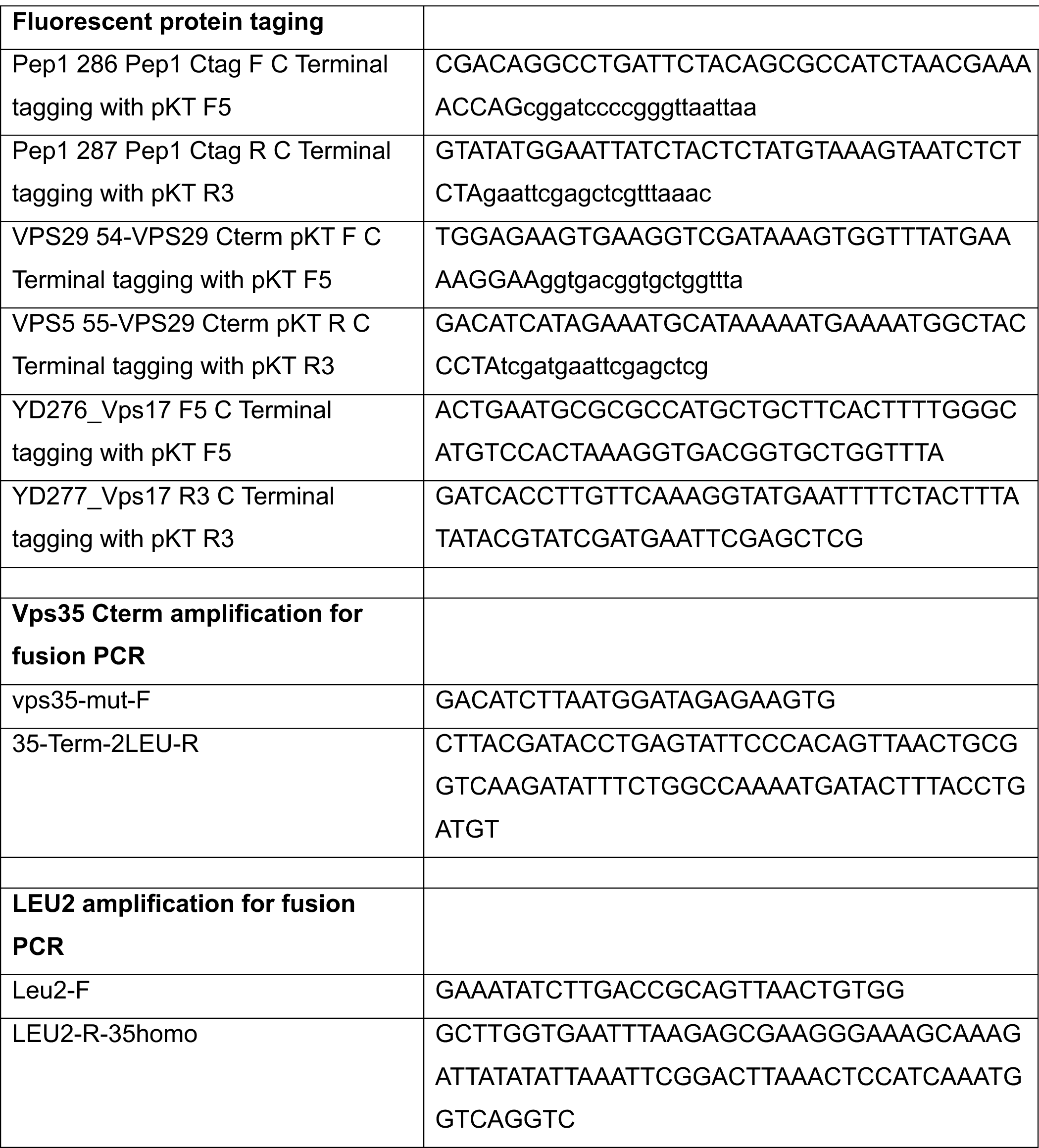
primers used in this study.

